# RAPIDASH: A tag-free enrichment of ribosome-associated proteins reveals compositional dynamics in embryonic tissues and stimulated macrophages

**DOI:** 10.1101/2023.12.07.570613

**Authors:** Teodorus Theo Susanto, Victoria Hung, Andrew G. Levine, Craig H. Kerr, Yongjin Yoo, Yuxiang Chen, Juan A. Oses-Prieto, Lisa Fromm, Kotaro Fujii, Marius Wernig, Alma L. Burlingame, Davide Ruggero, Maria Barna

## Abstract

Ribosomes are emerging as direct regulators of gene expression, with ribosome-associated proteins (RAPs) allowing ribosomes to modulate translational control. However, a lack of technologies to enrich RAPs across many sample types has prevented systematic analysis of RAP number, dynamics, and functions. Here, we have developed a label-free methodology called RAPIDASH to enrich ribosomes and RAPs from any sample. We applied RAPIDASH to mouse embryonic tissues and identified hundreds of potential RAPs, including DHX30 and LLPH, two forebrain RAPs important for neurodevelopment. We identified a critical role of LLPH in neural development that is linked to the translation of genes with long coding sequences. Finally, we characterized ribosome composition remodeling during immune activation and observed extensive changes post-stimulation. RAPIDASH has therefore enabled the discovery of RAPs ranging from those with neuroregulatory functions to those activated by immune stimuli, thereby providing critical insights into how ribosomes are remodeled.

## Introduction

Embryonic development is a highly regulated process that depends on the synthesis of the correct proteins at the correct time, place, and quantities in order to shape an organism. Translational control has been increasingly recognized as an important layer of regulation in gene expression during key stem cell fate decisions^1–3^, responses to stimuli^4–6^, and within developing embryos^7, 8^. Recently, the ribosome itself has emerged as a direct regulator of gene expression. One interesting point of regulation is the association of accessory proteins, such as RNA binding proteins, with the ribosome to exert specificity and regulation over mRNA translation. In particular, the ribosome has increased in size during eukaryotic evolution by virtue of expansion segments (ESs) that are inserted into ribosomal RNA (rRNA)^9^. Recent findings revealed hundreds of possible ribosome-associated proteins (RAPs) that may interact with the mammalian ribosome, possibly due to the expansion of rRNA, collectively known as the ribo-interactome^10^. A handful of these RAPs have been further functionally characterized, such as MetAP^11^, FMRP^12–14^, PKM^10, 15^, RBPMS^2^, UFL1^10, 16^, and CDK1^10, 17, 18^.

Key examples of RAPs include MetAP (also known as Map1 and Map2 in yeast) that directly binds to an expansion segment of the 28S rRNA, ES27L^11^ and is responsible for cleaving the initiator methionine of polypeptides as they are synthesized. Surprisingly, MetAP is important for translation fidelity^11^. FMRP binds directly to uL5 (Rpl11) and to neuronal mRNAs to inhibit their translation^12–14^. Pyruvate kinase (PKM), which catalyzes the final step of glycolysis to form pyruvate, is an RNA binding protein^19^ that directly interacts with ribosomes in mouse embryonic stem cells (mESCs)^10^. In mESCs, the dominant isoform is PKM2. Knockdown of PKM2 decreases the translational efficiency of mRNAs that are bound by PKM2, which indicates PKM2 is a translational activator for a subset of mRNAs^10^. RBPMS is an RNA binding protein that regulates the translation of mRNAs crucial for mesoderm differentiation. Knockdown of RBPMS leads to the inability to differentiate down the cardiac mesoderm lineage^2^. In addition to these proteins, post-translational modification enzymes interact with the ribosome to modify it. UFL1 is the sole known enzyme that deposits a post-translational modification known as ufmylation on proteins such as Rpl26, where the modification is associated with an increase in the stalling of ER translocon-associated ribosomes^16^. This suggests UFL1 helps regulate ribosome quality control, the ER stress response, and ER homeostasis. Lastly, cyclin-dependent kinase 1 (CDK1) phosphorylates Rpl12, which promotes translation of mitosis-related mRNAs^17^. Thus, RAPs connect the ribosome and translation to many important cellular functions.

Despite the importance of RAPs in diversifying translational control, there has been a lack of methods to rapidly and systematically identify RAPs in different tissue and cell types, which has stymied our ability to understand fundamental aspects of RAP biology. For example, how many RAPs are there, and do RAPs differ between cell types, tissues, and species? In the past, most work focused on identifying RAPs relied on fractionation of cellular contents according to density^20, 21^ or size^22^. However, these techniques are highly nonspecific. For example, endoplasmic reticulum (ER) microsomes have heterogeneous densities and thus are present throughout most fractions, including those containing ribosomes (Supplementary Figure 1A). More recent efforts have been much more specific to RAPs. For example, our lab used a genetic approach, which we refer to as Ribo-FLAG immunoprecipitation (IP), where the core RPs were endogenously tagged with FLAG epitope tag to enable the IP of ribosomes and RAPs for subsequent analysis by mass spectrometry (MS). This enabled the discovery of hundreds of RAPs in mouse embryonic stem cells^10^. Although Ribo-FLAG IP is exceptionally specific compared to methods that rely on fractionation, the reliance on genetic editing prevents us from easily implementing this strategy in many different sample types, such as tissues or patient samples. A chemical approach, active ribosome capture–mass spectrometry (ARC-MS), was recently reported as an orthogonal strategy to isolate ribosomes and RAPs^2^. Here, cells are pulsed with azidohomoalanine, a methionine analog that incorporates into the growing polypeptide chain in actively translating ribosomes. This provides a handle for subsequent Click chemistry with dibenzocyclooctyne beads to isolate ribosomes and RAPs for analysis by MS. Although this technique led to the identification of hundreds of RAPs in human embryonic stem cells and human embryonic stem cell-derived mesoderm^2^, its reliance on azidohomoalanine requires treating samples with a methionine analog and biases the findings towards ribosomes at the early steps of translation.

To bridge the gap in the field in systematically characterizing RAPs, here we develop a methodology that relies on the biophysical properties of the ribosome without genetic tagging or chemical modification, namely, the fact that it contains a core of ribosomal RNA. To do this, we turned to a chromatographic technique that had been used to isolate active ribosomes from bacteria^23^ and yeast^24^. This technique relies on a sulfhydryl-charged resin that enriches RNA. However, this protocol stripped all RAPs from the ribosomes. We therefore developed a new methodology based on this chromatography that preserves the interactions between RAPs and the ribosome for subsequent analysis by liquid chromatography-mass spectrometry (LC-MS). First, high density protein complexes are pelleted by sucrose cushion ultracentrifugation. These high density protein complexes are then subjected to chromatography with cysteine-charged resin to enrich RNA-containing, high density protein complexes, which are mostly ribosomes and their RAPs (Figure 1A). We called this methodology Ribosome-Associated Protein IDentification by Affinity to SulfHydryl-charged resin (RAPIDASH).

**Figure 1:**
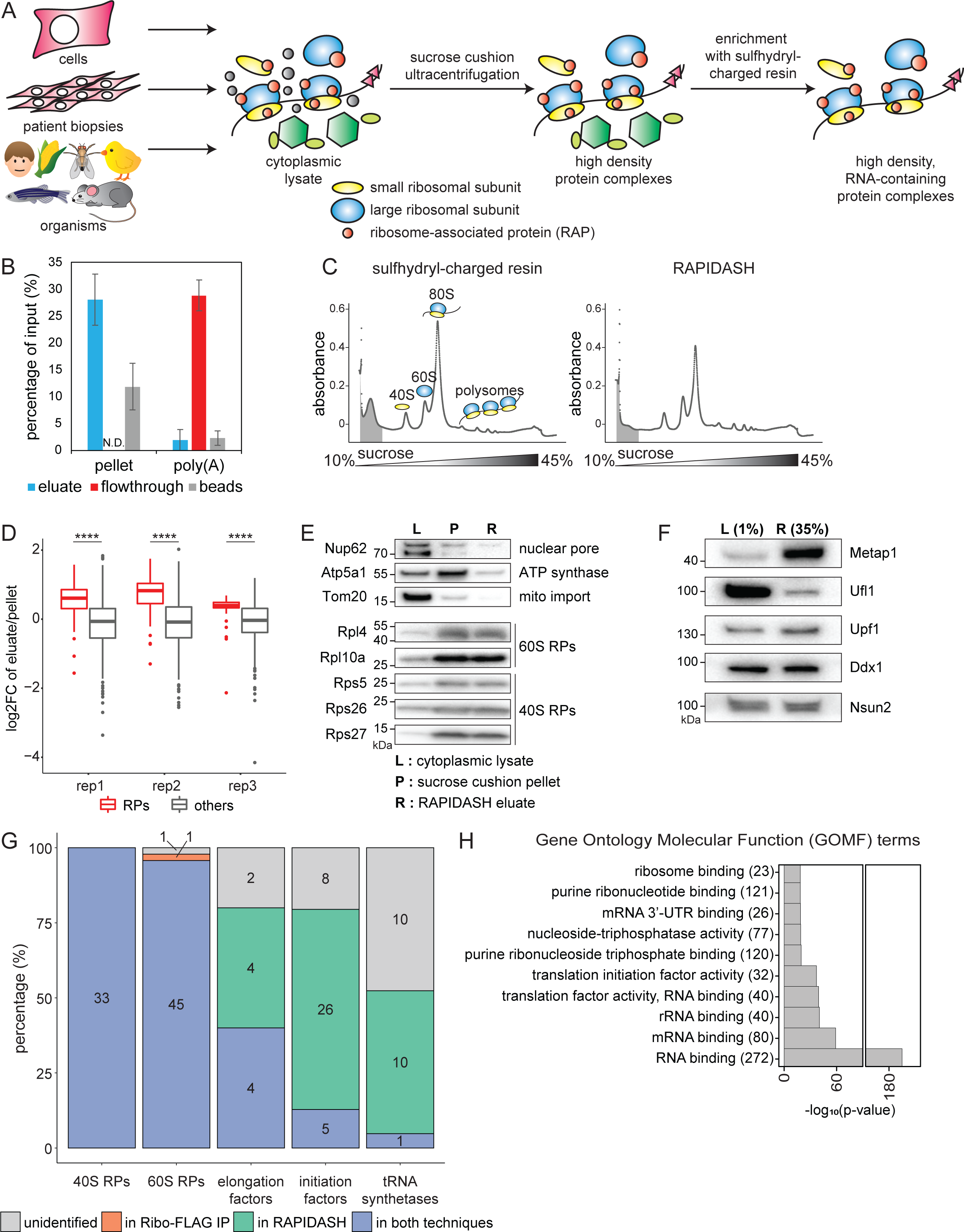
Characterization of the ribosome-associated protein identification by affinity to sulfhydryl-charged resin (RAPIDASH) method in E14 mouse embryonic stem cells (mESCs) (A) Schematic of the RAPIDASH protocol. This protocol can be applied to any biological sample, such as cells, patient biopsies, or organisms. Cytoplasmic lysates are subjected to sucrose cushion ultracentrifugation to enrich high density protein complexes. These then undergo enrichment for RNA-containing protein complexes by subjecting them to chromatography using sulfhydryl-charged resin. (B) Characterization of cysteine-charged sulfolink resin. Sucrose cushion pellet samples and poly(A) RNA isolated from mESCs were subjected to chromatography with cysteine-charged sulfolink resin. The percentage of RNA relative to input amount is plotted for the flowthrough, eluate, and bead-bound samples. N.D., not detected. Error bars are +/- standard error of the mean (SEM). (C) Characterization of RAPIDASH by sucrose gradient fractionation. Cytoplasmic lysate from E14 mouse embryonic stem cells (mESCs) was subjected to either enrichment with the sulfhydryl-charged resin or the entire RAPIDASH protocol. Each of these samples were fractionated on a sucrose density gradient to assess whether small ribonucleoproteins (gray) were depleted. (D) Boxplot of normalized log_2_ fold change (FC) of RAPIDASH eluate over sucrose cushion ultracentrifugation pellet tandem mass tag (TMT) mass spectrometry ratios from three biological replicates of mESCs. Ribosomal proteins (RPs; red) are significantly enriched over other proteins (gray) by RAPIDASH compared to sucrose cushion ultracentrifugation alone based on Welch’s t-test. p-values: rep1 = 1.24 × 10^-18^; rep2 = 4.39 × 10^-23^; rep3 = 8.30 × 10^-14^. (E) Western blotting of mESC sucrose cushion pellet and RAPIDASH eluate samples for components of non-ribosomal complexes to assess the specificity of RAPIDASH. Approximately equal amounts of RPs for sucrose cushion pellet and RAPIDASH eluate samples, as shown by SYPRO Ruby (Supplementary Figure 1E), were analyzed by western blotting for Nup62, Atp5a1, and Tom20, components of the nuclear pore complex, ATP synthase, and the translocase of the outer membrane (TOM) complex, respectively. Cytoplasmic lysate was included as an input control. (F) Western blot detection of known RAPs enriched by RAPIDASH. A representative blot with 1% of the mESC cytoplasmic lysate volume and 35% of the RAPIDASH eluate volume was probed for the known RAPs Metap1, Ufl1, Upf1, Ddx1, and Nsun2^10,11^. (G) Bar graph showing the percentage of translational machinery identified by Ribo-FLAG immunoprecipitation (IP)^10^ or RAPIDASH. Ribo-FLAG IP proteins are those that were identified by FLAG IP liquid chromatography-tandem mass spectrometry (LC-MS/MS) of endogenously FLAG-tagged Rpl36 or Rps17 in E14 mESCs^10^. Three biological replicates of RAPIDASH were performed for each of the same cell lines and analyzed by LC-MS/MS. Proteins identified with detectable peptide signal intensity in at least three out of the six RAPIDASH samples were compared against the Ribo-FLAG IP proteins. The percentage of 40S and 60S ribosomal proteins (RPs), translation elongation factors, translation initiation factors, and transfer RNA (tRNA) synthetases identified only in Ribo-FLAG IP (red), only in RAPIDASH (green), in both techniques (blue), or none (gray) are displayed. (H) Gene ontology (GO) term analysis of proteins identified by mESCs subjected to RAPIDASH. Proteins that were identified in RAPIDASH-enriched mESC samples were analyzed by Manteia^30^. The ten most significant GO molecular function (GOMF) terms whose minimum level was 4 are shown.

Herein, we have used RAPIDASH on mouse embryonic tissues including forebrain, limb, and liver and show that ribosomes can differ in RAP composition across tissue types. We focused on RAPs in the forebrain and characterized Dhx30, Elavl2, and LLPH as novel *bona fide* RAPs. We further characterized the role of LLPH in neurons, which revealed a critical role of LLPH in neural development and function. In addition, through ribosome profiling we strikingly found that LLPH has a selective role in translational control, in particular of long transcripts that may require more ribosome-directed control of mRNA translation. Furthermore, we have also applied RAPIDASH to characterize dynamic changes in ribosome composition at the level of RAPs in primary macrophages that are treated with different stimuli. This demonstrated the broad applicability of RAPIDASH to many sample types and to characterize how ribosome composition is remodeled in macrophages after stimulated bacterial and viral infections.

## Results

### Characterization of RAPIDASH in mouse embryonic stem cells (mESCs)

Here, we turned to a chromatographic technique that had been used to isolate active ribosomes from bacteria^23^ and yeast^24^ by binding of RNA moieties, likely rRNA, to a cysteine-charged resin and optimized it for the detection of RAPs. While the sulfhydryl-charged resin was used to isolate ribosomes, this protocol did not maintain interactions between the ribosomes and RAPs. In order to optimize the conditions to maximize binding and elution of mammalian ribosomes and to preserve interactions between ribosomes and RAPs, we made several changes to the protocol that was developed for yeast ribosomes^24^: (1) the order of operations was adjusted such that centrifugation was performed prior to chromatography with the resin, (2) ultracentrifugation speed was increased, but the time was decreased, and (3) the buffers were changed in order to maintain interactions with RAPs whose binding is salt sensitive. We isolated ribosomes using the sulfhydryl-charged resin according to the protocol for yeast ribosomes and compared the enriched proteins to those identified by our optimized RAPIDASH protocol using mass spectrometry to determine relative levels using TMT labeling. Out of the 201 proteins identified, only 51 proteins gave quantifiable signal in the reporter ion channel for the original chromatography protocol (Supplementary Table 1). Most of the quantified proteins were RPs. This number of proteins is far below what was identified by other methods used to characterize RAPs, which suggests the original chromatography protocol, while suitable to purify the core ribosome, cannot be used to comprehensively identify RAPs.

We then characterized the enrichment properties of the resin under our optimized conditions by performing the chromatography with either sucrose cushion pellet material or with poly(A)-enriched RNA. We found that sucrose cushion pellet material binds to and elutes from the resin. Importantly, this binding is selective as poly(A)-enriched RNA does not bind to the resin (Figure 1B). We also compared mESC samples subjected to cysteine-charged resin chromatography alone or to the entire RAPIDASH protocol by sucrose gradient fractionation to assess whether ribosomes are enriched over small ribonucleoproteins (RNPs). Sulfhydryl-charged resin chromatography enriches both small (area shaded in gray) and large RNPs, while the entire RAPIDASH protocol depletes the small RNPs but retains ribosomes (Figure 1C).

To assess if adding the cysteine-charged resin chromatography step allows for further enrichment of RPs compared to only performing sucrose cushion ultracentrifugation, mESC cytoplasmic lysates were subjected to either sucrose cushion ultracentrifugation alone or the entire RAPIDASH protocol. The proteins in each sample were digested to peptides and labeled with tandem mass tag (TMT) reagents according to Table 1 to allow for relative quantification by mass spectrometry between the samples. Samples were then pooled and then analyzed by liquid chromatography-tandem mass spectrometry (LC-MS/MS) (Supplementary Figure 1B). We detected between 823 to 874 proteins across all three replicates. Figure 1D shows that compared to sucrose cushion ultracentrifugation alone, RAPIDASH significantly enriches ribosomal proteins (RPs) over other proteins. Although there might be a slight bias for 60S (large subunit) RPs over 40S (small subunit) RPs, this is not significant across all three replicates (Supplementary Figure 1C).

**Table 1:** Information for samples prepared for relative quantification by TMT MS. Each TMT replicate constitutes a single injection of the mixed protein samples.

| **TMT replicate** | **protein sample** | **amount of protein (µg)** | **TMT tag** |
| --- | --- | --- | --- |
| RAPIDASH eluate vs. sucrose cushion pellet in mESCs, rep 1 | sucrose cushion pellet | 25 | 130 |
|  | RAPIDASH eluate | 25 | 131 |
| RAPIDASH eluate vs. sucrose cushion pellet in mESCs, rep 2 | sucrose cushion pellet | 25 | 130 |
|  | RAPIDASH eluate | 25 | 131 |
| RAPIDASH eluate vs. sucrose cushion pellet in mESCs, rep 3 | sucrose cushion pellet | 25 | 130 |
|  | RAPIDASH eluate | 25 | 131 |
| E12.5 mouse embryonic tissues, rep 1 | limbs | 15 | 126 |
|  | forebrain | 15 | 127 |
|  | liver | 15 | 128 |
| E12.5 mouse embryonic tissues, rep 2 | limbs | 25 | 126 |
|  | forebrain | 25 | 127 |
|  | liver | 25 | 128 |
| E12.5 mouse embryonic tissues, rep 3 | limbs | 13 | 126 |
|  | forebrain | 13 | 127 |
|  | liver | 13 | 128 |
| E12.5 mouse embryonic tissues, rep 4 | limbs | 25 | 126 |
|  | forebrain | 25 | 127 |
|  | liver | 25 | 128 |

We then asked whether non-ribosomal, large protein complexes are depleted after the entire RAPIDASH protocol compared to sucrose cushion ultracentrifugation alone. To answer this, we ranked the proteins by their average enrichment ratios of RAPIDASH compared to sucrose cushion ultracentrifugation across three replicates and asked which gene ontology cellular component (GOCC) terms are enriched for proteins that are in the bottom 10%. The most significantly enriched terms for these proteins were related to mitochondria, nucleus, cytoskeleton, and vesicles (Supplementary Figure 1D, Supplementary Table 2). To confirm the depletion of non-ribosomal, large complexes and structures by RAPIDASH, we performed western blot analysis of mESC sucrose cushion ultracentrifugation pellet versus RAPIDASH eluate samples for components of these complexes. As expected, when approximately equal amounts of RPs were loaded for each sample (Supplementary Figure 1E), the nuclear pore complex (Nup62), adenosine triphosphate (ATP) synthase (Atp5a1), and the translocase of the outer membrane (TOM) complex (Tom20) were depleted in the RAPIDASH sample compared to the sucrose cushion ultracentrifugation pellet sample (Figure 1E).

Subsequently, we assessed whether the RAPIDASH workflow can maintain interactions between ribosomes and their associated proteins. To do this, mESC cytoplasmic lysate was subjected to RAPIDASH and probed by western blotting for the presence of known RAPs. Ufl1^10, 16^, Upf1^25^, Ddx1^10^, Nsun2^10^, and Metap1^11^ were all present in the RAPIDASH samples (Figure 1F, Supplementary Figure 1F).

We next wanted to assess the sensitivity of RAPIDASH in capturing translation components. To do this, we examined the proportion of proteins in canonical translational machinery, that is, RPs, elongation factors, initiation factors, and the tRNA synthetase complex, that was identified by RAPIDASH. In addition, we compared our coverage with that of Ribo-FLAG IP, which is the most selective method to enrich ribosomes. We performed LC-MS/MS on RAPIDASH eluate derived from three biological replicates of Rps17-FLAG and Rpl36-FLAG mESCs each for a total of six samples. We detected 665 proteins present in at least 3 out of 6 replicates (Supplementary Table 3). Figure 1G shows the comparison of proteins in translational machinery that are identified by Ribo-FLAG IP using Byonic (v2.12.0)^26^ and analyzed by SAINT (v2.5.0)^27^ as described previously^10^, or in at least three out of six RAPIDASH samples identified using MaxQuant (v1.6.5.0)^28, 29^. RAPIDASH identified all of the RPs except for Uba52 (Rpl40) and Rpl41. Compared to Ribo-FLAG IP, RAPIDASH captured more translation elongation factors, initiation factors, and transfer RNA (tRNA) synthetases (Figure 1G), which suggests RAPIDASH has good coverage of RAPs. In addition, RAPIDASH detected 280 out of the 428 (65%) proteins identified by Ribo-FLAG IP^10^, which suggests RAPIDASH yields a high quality list of candidate RAPs. We also compared our mESC RAPIDASH data to the recent work of ARC-MS in human embryonic stem cells (hESCs)^2^. Out of the 665 proteins identified in RAPIDASH MS samples, there are 349 proteins that were also identified by ARC-MS in hESCs. This good overlap suggests that RAPIDASH is finding many RAPs identified by ARC-MS. Since ARC-MS is disposed towards finding proteins that are involved in the early steps of translation, the non-overlapping proteins may be those that are RAPs that are more involved in other steps of translation, such as elongation or termination. Other non-overlapping proteins may be explained by proteins that are RAPs in mice but not humans.

When we performed gene ontology analysis using Manteia^30^ on RAPIDASH MS samples from mESCs, most of the ten most significant gene ontology molecular function terms are those for RNA binding, either to mRNA or ribosomal RNA, or those terms dealing with translation or binding to the ribosome (Figure 1H). Together, these results suggest RAPIDASH is a method with good specificity and coverage that can identify candidate RAPs.

### Application of RAPIDASH to the developing mouse forebrain identifies Dhx30 as a RAP

Given the ability of RAPIDASH to preferentially isolate ribosomes in a clean, untagged manner, we next sought to identify RAPs present in the developing mouse forebrain tissue. We performed RAPIDASH on E12.5 forebrains followed by LC-MS/MS. This analysis yielded 594 protein identifications present in 2 out of 3 replicates (Supplementary Table 4). Gene ontology analysis^30^ of the data showed significantly enriched gene ontology molecular function terms such as RNA binding and mRNA binding (Figure 2A), suggesting that proteins identified are bona fide RAPs with potential forebrain-specific functions.

**Figure 2:**
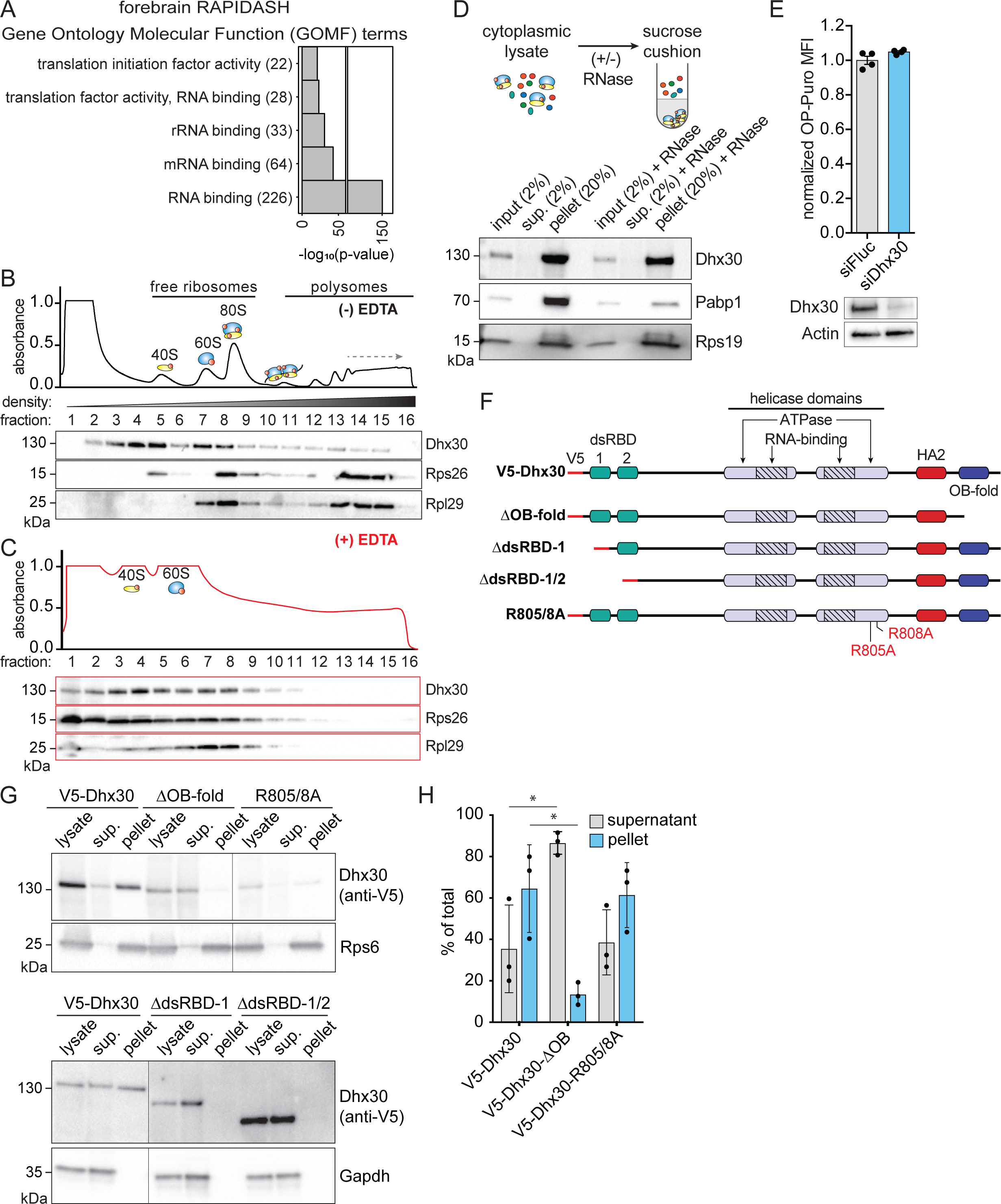
Characterization of Dhx30 as a bona-fide mRNA-independent RAP. (A) Analysis of GO terms in forebrain MS data. RAPIDASH was performed on E12.5 mouse forebrain samples. These samples were analyzed by LC-MS/MS. The resulting proteins were analyzed by Manteia^30^ for GOMF terms level 4 or higher. The top 5 GOMF terms are shown. (B) Dhx30 co-fractionates with ribosomes, specifically the 40S fractions. Sucrose gradient fractionation was performed on E14 mESCs. The proteins from each fraction were precipitated and analyzed by western blotting. Dhx30 does not co-fractionate with the free fraction; instead, it largely co-fractionates with the 40S and 80S fractions. Rps26 and Rpl29 are shown as controls for small and large subunits, respectively. (C) E14 mESCs were treated with EDTA and subjected to sucrose gradient fractionation followed by western blotting as in (B). (D) Dhx30 still associates with the pellet fraction after RNase treatment. Top: illustration of sucrose cushion and RNase A treatment of E14 mESC cytoplasmic lysate. Bottom: western blot of control and RNase A-treated E14 mESC subjected to sucrose cushion ultracentrifugation. Pabp1 is a positive control for mRNA-dependent association with ribosomes. Rps19 is a positive control for ribosomes. Sup., supernatant. (E) Knockdown of Dhx30 by siRNA in mESC does not affect global protein synthesis. Protein synthesis was measured by O-propargyl-puromycin (OP-Puro) incorporation into the nascent proteome. Incorporated OP-Puro was fluorescently labeled using a click chemistry reaction. Translation activity was then measured using flow cytometry. Top: there is no change in OP-Puro median fluorescence intensity (MFI) between siFluc and siDhx30 (n=4). Bottom: western blotting of Dhx30 shows that siRNA knockdown is successful. (F) Illustration of Dhx30 constructs for transient transfection in E14 mESCs. Oligosaccharide binding (OB) fold domain; and double-stranded RNA-binding domains (dsRBDs). (G) Top: western blots of E14 mESCs transfected with V5-Dhx30, ΔOB-fold, and R805/8A constructs and subjected to sucrose cushion ultracentrifugation. Bottom: similar samples, but instead transfected with V5-Dhx30, ΔdsRBD-1, and ΔdsRBD-1/2 constructs. Loss of either OB-fold, dsRBD-1, or dsRBD-2 results in the loss of Dhx30 association with the ribosome. However, Dhx30 loss of function due to loss of helicase activity in R805/8A mutant does not affect its association with the ribosome. (H) Quantification of Dhx30 western blots shows a significant shift from pellet to supernatant in ΔOB-fold transiently transfected E14 mESCs but not R805/8A transfected E14 mESCs.

To identify hits for further functional characterization, we separated the identified proteins into two classes: (I) proteins that are known to regulate translation but are not known ribosome binders, and (II) proteins known to bind to the ribosome that are not known to regulate translation. Following this schema, we identified Dhx30 as a paradigm Class I protein. Dhx30 belongs to the DExH class of RNA helicases. DExD/H-box RNA helicases are highly conserved enzymes that use the energy of ATP to remodel RNA secondary structure and are vital for regulating various aspects of the RNA life cycle that when perturbed can lead to disease^31^.

Biallelic loss of *Dhx30* leads to embryonic lethality in mice exhibiting distinct early neural defects^32^. Strikingly, mutations mapped to the helicase and RNA-binding domains of human DHX30 have been directly identified in pediatric neurodevelopmental disease characterized by severely delayed psychomotor development and muscular hypotonia at early infancy resulting in gait abnormalities or inability to walk, speech impairment, and severe intellectual disability^33, 34^. Dhx30 has been implicated in cytoplasmic translational regulation and is suggested to bind to mRNAs that carry a 3’ UTR CG-rich motif mediating p53-dependent death (CGPD-motif) and decrease their translation^35^.

To test whether Dhx30 is a direct ribosome binder, we first performed sucrose gradient fractionation on mouse embryonic stem cells (Figure 2B), where cytoplasmic lysate is separated according to density. Each fraction was precipitated and probed for ribosomal protein markers or for Dhx30. We observed that Dhx30 co-fractionates with ribosomal subunits, mature 80S ribosomes, and polysomes, as marked by the ribosomal proteins Rps26 and Rpl29 (Figure 2B). Furthermore, EDTA-mediated dissociation of mature 80S ribosomes and polysomes disrupts the Dhx30 co-fractionation profile along the gradient (Figure 2C). We next tested whether the interaction between Dhx30 and the ribosome is dependent on mRNA. To do this, we split cytoplasmic lysate into two samples and treated one with RNase A to digest mRNA. Each sample was then subjected to ultracentrifugation over a sucrose cushion to enrich high density protein complexes, including ribosomes and their associated proteins. Dhx30 showed comparable signal in the pellets derived from the untreated and RNase-treated cytoplasmic samples, while a known mRNA-binding protein, PABPC, serving as a control demonstrated depletion in the RNase A-treated samples (Figure 2D). This indicates Dhx30 interaction with the ribosome is mRNA-independent. We next asked if Dhx30 regulates translation globally like other RNA helicases such as eIF4A or DHX29. Knock down of Dhx30 with siRNA did not significantly change the global protein synthesis rate in mESCs compared to a non-specific siRNA control (Figure 2E). These data support prior literature describing Dhx30 mediation of translation of specific subsets of mRNAs.

As Dhx30 is a protein with multiple functional domains, we next asked which domains are responsible for the binding of Dhx30 to the ribosome. Dhx30 has two OB-fold domains, two dsRBD domains, a core of two helicase domains, and a helicase-associated domain (HA2) (Figure 2F). To test which domain is involved in ribosome-interactions, we generated a variety of V5-tagged Dhx30 mutants that harbor either human disease-relevant mutations which inactivate helicase activity (R805/8A)^33^ or domain truncations (Figure 2F). Each mutant was then transfected into mESCs and cell lysates separated by ultracentrifugation over a sucrose cushion. We observed an enrichment of both wild-type Dhx30 and the R805/808A double mutant in the pellet compared to the supernatant suggesting that helicase activity is not required for ribosome binding. Importantly, we observed almost complete loss of ribosome association in the ΔOB-fold and ΔdsRBD (Figures 2G and 2H). Taken together, these data suggest that Dhx30 binds the ribosome through its OB-fold and dsRBD domains. These findings thereby identify a new bona fide RAP from our RAPIDASH results and distinguish the domains of Dhx30 that bind to the ribosome.

### LLPH as a RAP important for neurodevelopment

Our forebrain data also yielded a paradigm Class II protein (those known to bind to the ribosome but are not known to regulate translation) called LAPS18-Like Protein Homolog (LLPH). The homolog, Learning-Associated Protein of Slug with 18 kDa (LAPS18), was first functionally characterized in *Limax marginatus*, where it was suggested to be a secreted protein important for long term memory formation^36^. In contrast, work studying the *Aplysia kurodai* homolog *Aplysia* LAPS18-like protein (ApLLP) described ApLLP as a nuclear protein enriched in the nucleolus that is important for the switch to long term facilitation^37, 38^. In mice, LLPH is highly expressed in the developing mouse brain, but its expression wanes once the pup is born. Further work studying mouse LLPH characterized it as a cell-permeable protein that is important for neural development: knockdown of LLPH in cultured hippocampal neurons impairs dendritic growth and results in shorter neurites^39^. Our RAPIDASH data suggest that LLPH may link the ribosome to a critical role in neural development and function.

The N-terminus of LLPH is highly conserved and forms a helix, while the remainder of the protein is disordered. Recently, cryo-electron microscopy (cryo-EM) structures of the pre-60S particle in yeast^40^ and humans^41^ have captured the N-terminus of LLPH binding to the ribosome at the base of sarcin-ricin loop (Figure 3A). This binding site is suggestive of LLPH’s potential role in regulating translation, as the sarcin-ricin loop is a highly conserved sequence critical for GTP hydrolysis of EF1A and EEF2, which enables proper translocation during protein synthesis^42, 43^. Therefore, LLPH may be important for regulating translation elongation.

**Figure 3:**
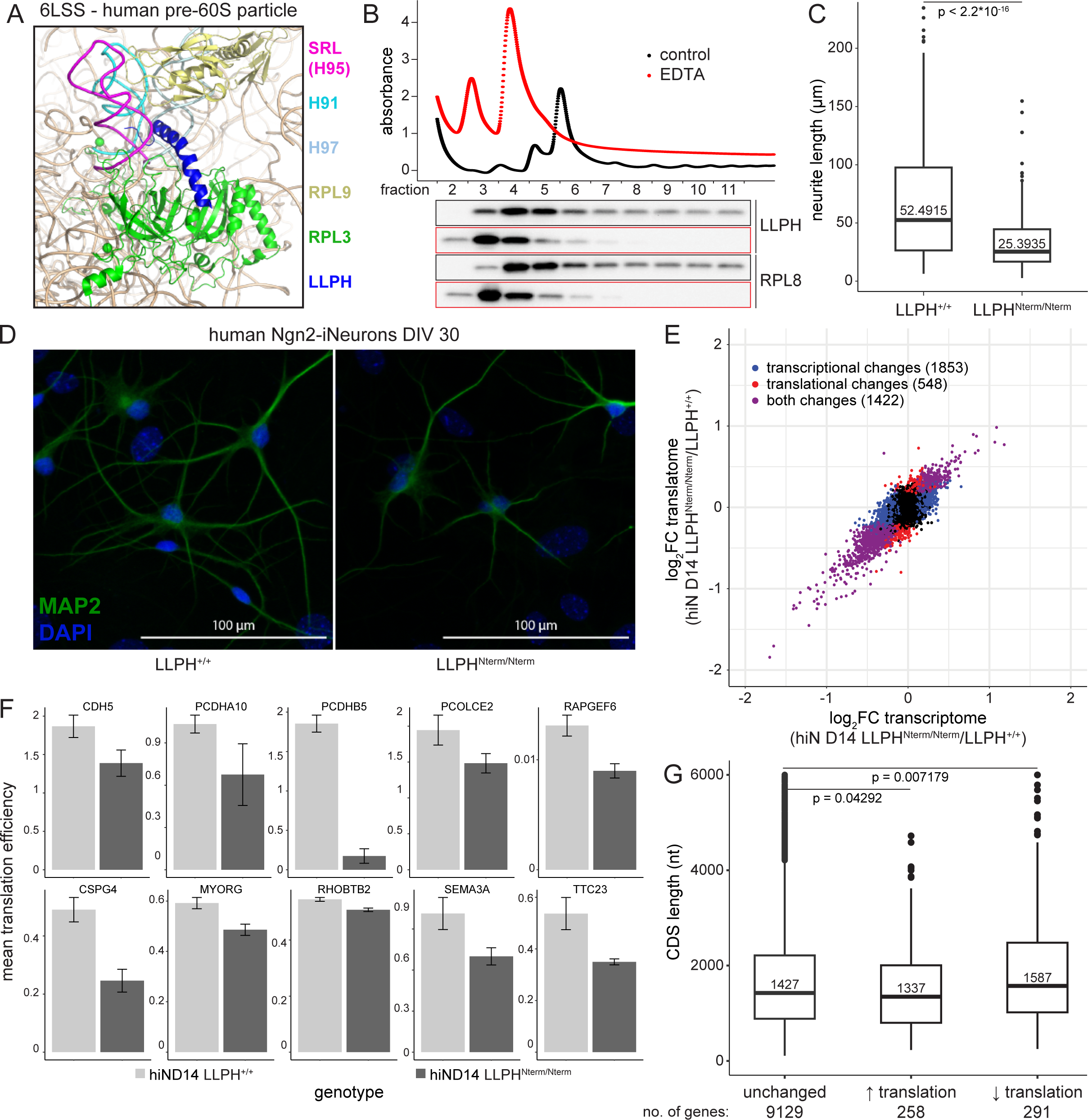
LLPH is a novel RAP with a role in neurodevelopment. (A) LLPH binding location on the ribosome. Previously published cryo-EM data show LLPH binds near the sarcin-ricin loop of the human pre-60S particle, a highly conserved region in the ribosome critical for elongation (Protein Data Bank (PDB) ID: 6LSS). (B) LLPH cofractionates with ribosomes, specifically 60S fractions. Sucrose gradient fractionation was performed with (red) or without (black) EDTA in P493-6 cells where LLPH is highly expressed. The proteins from each fraction were precipitated and analyzed by western blotting. Rpl8 is a control for the large subunit. (C) Measurement of traced primary neurite lengths of individual LLPH^+/+^ and LLPH^Nterm/Nterm^ human Ngn2-induced neurons (hiNs) at days in vitro (DIV) 30. LLPH^Nterm/Nterm^ hiNs have shorter neurites, which may hint at neurodevelopmental defects. (D) Representative fluorescence images of fixed DIV 30 wild-type and LLPH^Nterm/Nterm^ hiNs. Wild-type and LLPH^Nterm/Nterm^ hiNs were fixed and stained with a primary antibody against MAP2 and 4’,6-diamidino-2-phenylindole) (DAPI). (E) Comparison of Ribo-seq and RNA-seq data for DIV 14 LLPH^+/+^ and LLPH^Nterm/Nterm^ hiNs (n = 3 each). Blue genes are those that significantly change (Benjamini-Hochberg procedure (FDR) < 0.1 and absolute fold change (FC) ≥ 2) in mRNA abundance only; red genes are those that change in ribosome occupancy only; purple genes are those that change in mRNA abundance and ribosome occupancy. (F) Representative genes with lower mean translation efficiency differences in LLPH^Nterm/Nterm^ vs. LLPH^+/+h^ hiNs. Top: representative genes with downregulated translation efficiency in LLPH^Nterm/Nterm^ compared to LLPH^+/+^ that are involved in building the extracellular matrix. Bottom: representative genes known to be linked to neurodevelopmental defects related to growth cone defects or mRNA transport. (G) Genes downregulated for translation tend to have longer coding sequences (CDSs) than those that are translationally unchanged (Mann-Whitney U test, p = 0.0072). For clarity, only genes with CDS lengths shorter than 6000 are displayed. The median CDS length in each condition is displayed inside each boxplot. The full plot is shown in Supplementary Figure 2F.

First, we wanted to confirm LLPH is a RAP by performing a sucrose gradient fractionation experiment. Because LLPH is easier to detect in human samples compared to mouse samples, we used P493-6 lymphoblastoid cells, which express LLPH at a high level. P493-6 cell lysate was fractionated, and the fractions were probed by western blotting for the presence of LLPH (Figure 3B, Supplementary Figure 2D). This showed LLPH associates mainly with the 60S subunit but is also present in mature ribosomes and polysomes. Treatment of the lysate with EDTA and RNase A prior to fractionation to dissociate ribosomes into their subunits shows that LLPH co-migrates with the 60S subunit (Figure 3B, Supplementary Figure 2D).

To extend these data to another human cell line and to identify possible binding partners of LLPH-containing ribosomes, we established an A549 cell line that expresses LLPH-Flag when treated with doxycycline. Immunoprecipitation of LLPH-Flag from these cells following doxycycline treatment and analysis by mass spectrometry revealed LLPH binds to the ribosome. Additional binding partners, which include the elongation factor EEF2; the RAPs PA2G4, SERBP1, CCDC124; the RTRAF-RTCB-DDX1-FAM98B complex; and the BTF3-NACA complex, hint at a possible role in translational control (Supplementary Figure 2A). For example, EEF2 is involved with ribosome translocation^44^. PA2G4 is an RNA-binding protein that binds near the exit tunnel on the 60S subunit and is involved in growth regulation^45–47^. SERBP1 and CCDC124 bind to hibernating ribosomes and help the recovery of translation^48^. The RTRAF-RTCB-DDX1-FAM98B complex binds to the cap of the mRNA and increases the rate of translation^49^. The BTF3-NACA binds to the nascent polypeptide and prevents non-endoplasmic reticulum proteins from being trafficked to the endoplasmic reticulum^50, 51^. These data suggest that LLPH may participate in different steps of translational control.

Next, we wanted to directly characterize LLPH’s role as a RAP in neurons. We established H1-human embryonic stem cell (H1-hESC) lines that could be differentiated into neurons. CRISPR-Cas9 was used to edit LLPH so that only the N-terminal 24 amino acids of LLPH, plus five extra amino acids that were the result of the genomic editing, were expressed (LLPH^Nterm/Nterm^) (Supplementary Figure 2B). Importantly, LLPH^Nterm/Nterm^ hESCs are indistinguishable from wild-type hESCs in terms of growth (Supplementary Figure 2D). We differentiated these hESCs into induced neurons (hiNs) by transducing hESCs with lentivirus to express neuroligin 2. To assess whether LLPH is expressed in hiNs and whether its interaction with the ribosome is RNA-dependent, we harvested days in vitro (DIV) 5 hiNs, lysed the cells, treated the lysate with or without RNase A, and performed sucrose gradient fractionation. The proteins in the fractions were analyzed by western blotting. This revealed that LLPH is present in DIV 5 hINs and binds to the ribosome in an RNase-independent manner (Supplementary Figure 2E). The morphology of the hiNs was assessed on days in vitro (DIV) 30. We observed that LLPH^Nterm/Nterm^ neurons form shorter neurites with fewer secondary and tertiary neurites compared to the wild-type control neurons (Figures 3C and 3D). Thus, the LLPH^Nterm/Nterm^ mutant phenocopies LLPH knockdown in cultured hippocampal neurons.

We next asked if there was a difference in translation regulation that could lead to the neural phenotype in LLPH^Nterm/Nterm^ neurons. We wanted to probe translation at an early stage during the differentiation to determine whether a difference in translation preceded the morphological differences. To do this, we performed ribosome profiling in DIV 14 wild-type and LLPH^Nterm/Nterm^ hiNs. We observed changes at the transcriptional and translational level for the LLPH^Nterm/Nterm^ mutants compared to wild-type cells (Figure 3E, Supplementary Table 5). Genes that were translationally downregulated in LLPH^Nterm/Nterm^ mutant hiNs included those involved with the extracellular matrix (ECM); such as a cadherin and two protocadherins and CDH5, PCDHA10, and PCDH5B; procollagen C-endopeptidase enhancer 2 (PCOLCE2)^52^; and the guanine exchange factor RAPGEF6, which has been linked to neuritogenesis^53^ (Figure 3F). This is striking because the extracellular matrix is important for many aspects of neurodevelopment^54, 55^. In addition, several non-ECM genes important for neurodevelopment are also translationally downregulated in the LLPH^Nterm/Nterm^ mutant cell compared to wild-type hINs, such as CSPG4^56^, MYORG^57^, RHOBTB2^58, 59^, SEMA3A^60^, and TTC23^61, 62^ (Figure 3F). These may help explain why LLPH^Nterm/Nterm^ hiNs fail to produce neurites of the same length as wild-type hiNs.

Finally, because the binding site of LLPH on the ribosome suggests a role in regulating elongation, we asked whether genes that are translationally downregulated in LLPH^Nterm/Nterm^ hiNs have any correlation with length because elongation rate is known to be negatively correlated with coding sequence length^63^. We strikingly found that genes having longer coding sequences are enriched compared to those that show no change in translation when comparing LLPH^Nterm/Nterm^ and wild-type LLPH hiNs, which suggests LLPH may have a role in promoting elongation (Figure 3G, Supplementary Figure 2F). This is intriguing in the context of neurons because neuronal genes can have long coding sequences. Since coding sequence length also negatively correlates with translation fidelity^64^, regulation of elongation may be crucial to faithfully translate neuronal genes. Thus, LLPH may allow the translation of genes that are important for neural development.

### Identification and characterization of tissue-specific RAPs in developing mouse embryo

Having confirmed that RAPIDASH can discover RAPs in tissue samples, we next asked whether RAPs can change across embryonic tissues. We microdissected the limbs, forebrain, and liver of E12.5 FVB/NJ mouse embryos and prepared four biological replicates of each tissue. Each tissue sample was separately subjected to RAPIDASH, digested to peptides, and labeled with TMT reagents. The peptides from each tissue sample were mixed and then analyzed by LC-MS/MS (Figure 4A).

**Figure 4:**
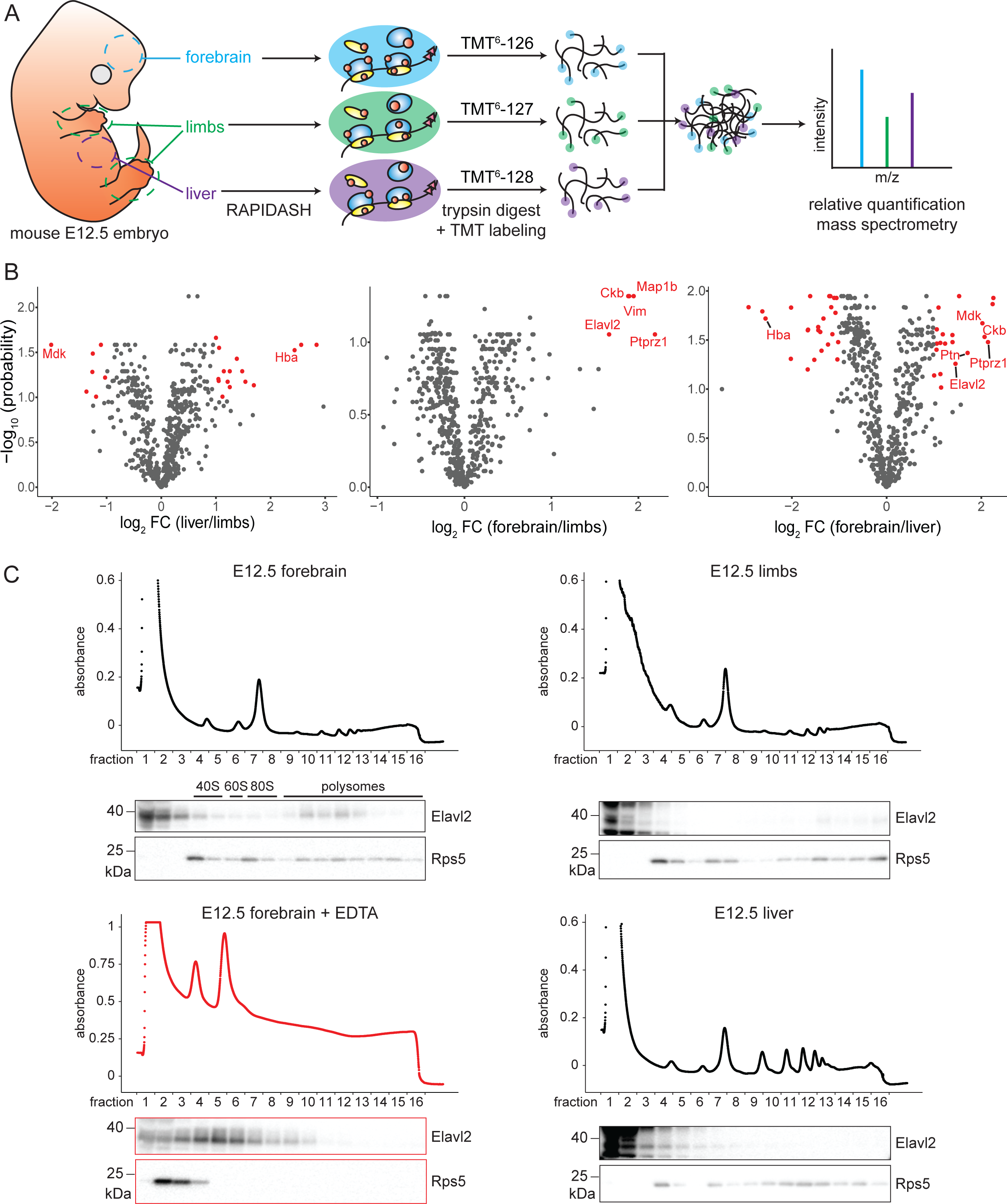
Characterization of tissue-specific RAPs in the E12.5 mouse embryo. (A) Schematic of the strategy to identify and quantify tissue-specific RAPs by tandem mass tag (TMT) mass spectrometry. Forebrain, limbs, and liver tissues were microdissected from E12.5 FVB/NJ mouse embryos and subjected to RAPIDASH. The enriched proteins were digested to peptides, which were labeled with TMT reagents to allow for relative quantification by LC-MS/MS. Four biological replicates were performed. (B) Volcano plots showing RAPs that are significantly enriched in one tissue compared to another. Putative RAPs identified in: liver to limbs (left), forebrain to limbs (center), forebrain to liver (right) are shown using volcano plots graphing -log_10_(p-value) against log_2_(FC). Proteins present in at least three out of four biological replicates with |log_2_FC| ≥ 1 and false discovery rate (FDR) < 0.10 are defined as significantly differentially enriched (red). (C) Elavl2 is a forebrain-enriched RAP. Forebrain, limb, and liver tissues from E12.5 mouse embryos were separated by sucrose gradient fractionation. An additional sample of forebrain tissue treated with EDTA as a control was also subjected to sucrose gradient fractonation. The protein from each fraction was precipitated and analyzed by western blotting for the presence of Elavl2 or Rps5, which served as a marker for the ribosome.

Each of the four biological replicates had between 531 to 641 proteins (Supplementary Table 6). We required proteins to be detected and quantified in at least three out of four replicates for our final analysis. Proteins that had FC ≥ 2 or FDR < 0.1 when comparing one tissue to another were considered enriched in a particular tissue. When these data were graphed on volcano plots to identify putative tissue-enriched RAPs, comparing forebrain and limbs samples to liver samples yielded more proteins that passed the significance cutoffs than for forebrain versus limbs samples (Figure 4B). This might be due to the more homogenous cell composition in liver tissue compared to the other tissues, which highlights the inter-tissue differences. Hemoglobin subunit alpha (Hba) was shown to be enriched in the liver as a RAP compared to the other two tissues, which hints for the ribosome’s role in heme bioavailability regulation as described previously^65^. Hkdc1, a kinase that phosphorylates hexose to hexose-6-phosphate in glycolysis, is enriched as a RAP in the liver relative to the other two tissues. This finding is reminiscent of how Pkm, another metabolic kinase, was shown to directly interact with ER ribosomes and regulate translation of specific mRNAs^10^.

The forebrain and limbs had very few RAPs that were differentially enriched; all five RAPs that were differentially enriched between these two tissues were enriched in the forebrain relative to the limbs. Creatine kinase b (Ckb), a kinase important for energy transduction in energy intensive tissues (e.g. muscle, heart, and brain)^66^, is enriched as a RAP in the forebrain over both the liver and limbs. Vimentin is an intermediate filament protein that is crucial for spatially organizing organelles within cells, and it has been implicated in cell adhesion^67, 68^, cell migration^68^, and neural development. It has also been observed to bind to ribosomes *in vitro*, which suggests that vimentin may help anchor ribosomes in the complex spatial architecture of neurons^69^.

Strikingly, we also identified the secreted growth factors midkine (Mdk) and pleiotrophin (Ptn), as putative RAPs across all three tissues, although they were both enriched in the forebrain relative to the liver. (Figure 4B). Mdk and Ptn have many roles ascribed to them, some of which are shared, such as cell differentiation, inflammation, cancer, and development^70, 71^. A brain-specific transmembrane protein tyrosine phosphatase Ptprz1 is known as their receptor^72^, which we also found to be a forebrain-enriched RAP. All three of these proteins are involved in neural development; therefore, finding them enriched in the forebrain relative to the liver suggests that their binding to the ribosome may be important for this function. It is tempting to speculate that the ribosome is a substrate for Ptprz1, and the binding of the ligands Mdk or Ptn may modulate phosphatase activity on the ribosome and initiate a translational program that is important for neural development.

Finally, Elavl2 is an RNA-binding protein that was found as a forebrain-enriched RAP compared to both liver and limbs. Elavl2 binds to the 3’ untranslated regions (UTRs) of selected mRNAs^73^. Elavl2 is a paradigm class I protein; although it is an RNA-binding protein that has been suggested to be a translation repressor in ovaries^74^, it is possible that in these embryonic tissues, Elavl2 is also a direct ribosome binder. We performed a sucrose gradient fractionation experiment on embryonic forebrain, limbs, and liver tissue lysate that was treated with EDTA or left untreated, and confirmed Elavl2 is a RAP in tissues (Figure 4C). Furthermore, Elavl2 is more enriched in the ribosome fractions in the forebrain compared to other tissues, suggesting its role on the ribosome may be important for brain development. Taken together, RAPIDASH has revealed hundreds of potential RAPs in embryonic tissues that may play a role in tissue-specific functions.

### RAPIDASH reveals dynamic ribosome composition remodeling during macrophage stimulation

Finally, we leveraged RAPIDASH to understand how an acute stimulus could temporally remodel ribosome complexes within a given cell type. Among cells in the body, macrophages play particularly diverse roles in organismal homeostasis, including in clearance of apoptotic debris, iron recycling, and wound healing^75^. Macrophages are also essential for host defense, playing critical roles in immune response initiation, amplification, and resolution. Mechanistically, macrophages sense the presence of bacterial and viral pathogens via germline-encoded pattern recognition receptors, including the family of Toll-like receptors (TLRs), which each recognize a distinct ‘non-self’ structure^76^. For example, TLR4 senses invasion by Gram-negative bacteria via binding to their outer cell membrane component lipopolysaccharide (LPS). At the same time, TLR3 alerts the macrophage to potential viral infection via binding to double-stranded RNA. Each TLR, once activated by its ligand, induces a distinct transcriptional program tailored to defend the host against the particular pathogen detected (e.g. proinflammatory cytokine production downstream of TLR4 and type I interferon production downstream of TLR3.) We hypothesized that activation of TLRs in macrophages might induce similarly distinct RAP-ribosome interactions to create an unappreciated ribosomal composition that might be tailored to meet the needs of each challenge.

We first established that RAPIDASH could effectively isolate highly enriched ribosomes from unstimulated primary murine bone marrow-derived macrophages (BMDMs). By mass spectrometry, we identified a total of 1,237 proteins with at least one peptide in each of three replicates (Supplementary Table 7). As expected, GO term analysis showed strong enrichment for ribosome and translation-related processes (Supplementary Figure 3A). Ribosomal proteins were highly enriched: 31 of 33 small (94%) and 41 of 44 large (93%) ribosomal subunit proteins detected were present in the top 150 most abundant proteins. Consistent with efficient isolation of intact complexes, the top 150 proteins also included subunits of the translation initiation factors eIF2, eIF3, and eIF5; the elongation factor EEF2; and the known RAP PA2G4.

We next used TMT-MS to determine whether a specific challenge would remodel the composition of ribosome complexes. The macrophage response to infection by viruses—which seek to co-opt host ribosomes for virion replication—involves a well-characterized reorganization of the translational machinery via phosphorylation of translation initiation factors^77, 78^. We reasoned that viral challenge might also induce particularly robust RAP-ribosome interactions as part of the antiviral response. Therefore, we isolated ribosome complexes from BMDMs 0, 2, 6, 12, and 24 hours post-activation by the TLR3 agonist and double-stranded RNA (dsRNA) mimic polyinosinic-polycytidylic acid or poly(I:C). Indeed, we found that ribosome complexes were gradually remodeled over time (Figure 5A and B, Supplementary Tables 8 and 9). By 24 hours post-stimulation, 395 proteins were enriched 2-fold, and 59 proteins were de-enriched 2-fold compared to ribosome complexes in resting BMDMs. Confirming the ability of RAPIDASH to detect RAPs, a number of the most highly induced RAPs were proteins known to interact with the translational machinery during viral infection. These included the IFN-induced protein with tetratricopeptide repeats (IFIT) proteins IFIT1, IFIT2, and IFIT3 which inhibit translation of viral transcripts in part through physical interaction with the 43S pre-initiation complex^79^; the ubiquitin-like protein ISG15, which is co-translationally ligated onto viral proteins thus interfering with their function^80^; and the E3 ubiquitin ligase for ISG15, Herc6^81, 82^. We also identified known RAPs that are co-opted by viruses to promote their pathogenesis, including the deubiquitylase USP15, which associates with polysomes and may stabilize newly-synthesized (viral) proteins^83, 84^; and the prolyl hydroxylase P4HA1, which has been demonstrated to co-translationally modify proline residues of flavivirus proteins^85^. Among the 59 proteins ‘ejected’ from ribosome complexes upon TLR3 stimulation, we identified PDCD4, a known ribosome-interacting translational inhibitor^86^. We also identified HMGB1, HMGB2, and HMGB3 proteins; HMGB2 in particular has recently been implicated in ribosome biogenesis^87^ and has been identified as a putative RAP^88^. The function of this family of proteins on resting ribosome complexes remains to be explored.

**Figure 5.**
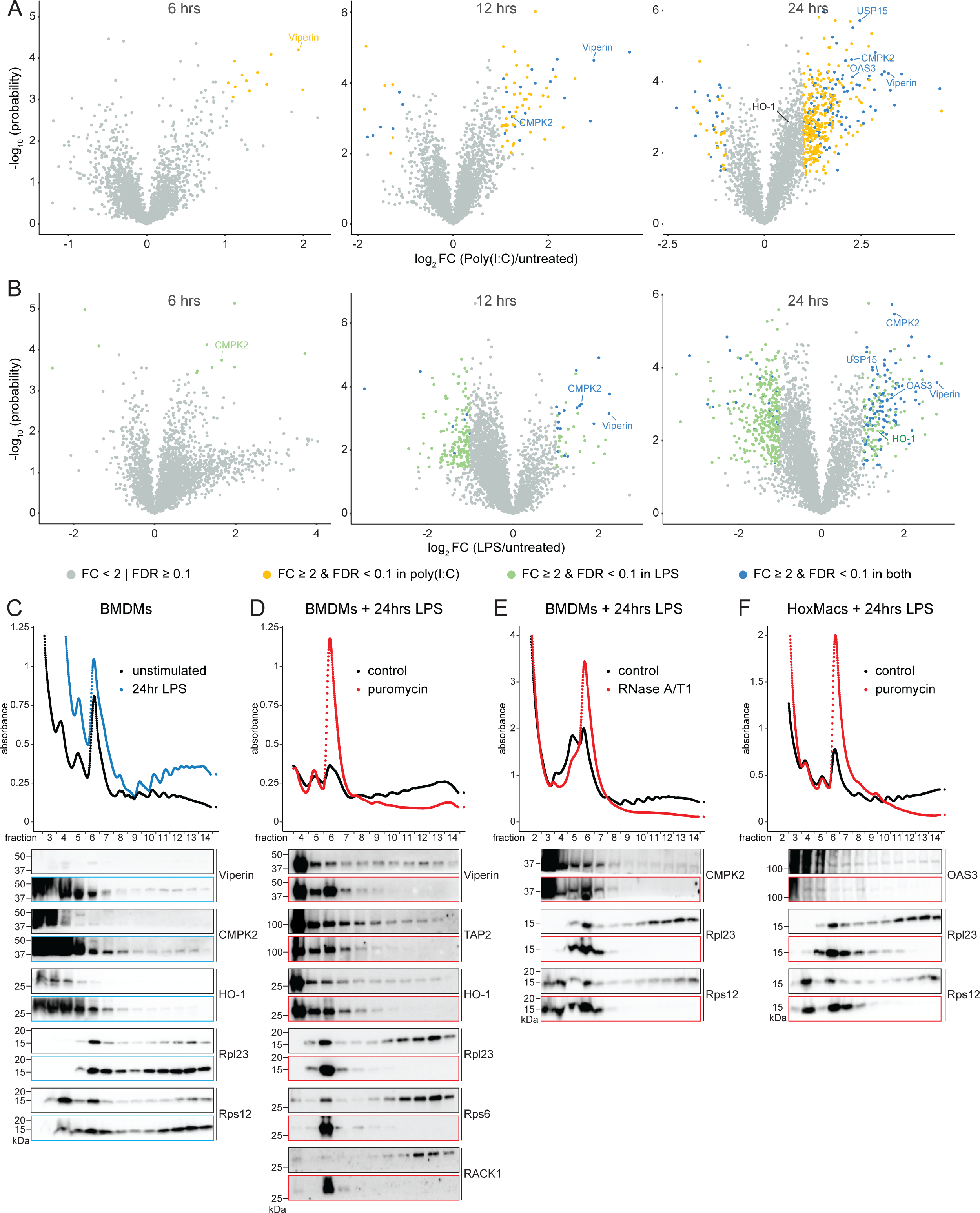
RAPIDASH identifies novel RAPs in macrophages following TLR stimulation. (A-B) Murine bone marrow derived macrophages (BMDMs) were stimulated with lipopolysaccharide (LPS) (A) or polyinosinic-polycytidylic acid (poly(I:C)) (B) for 6, 12, or 24 hours prior to isolation of ribosome complexes for TMT-MS analysis. X-axes show the log_2_ FC of ribosome complex composition in activated versus unstimulated macrophages at 6 hours (left panels), 12 hours (middle panels), and 24 hours (right panels). (C-F) Lysates from unstimulated or LPS-stimulated BMDMs (C-E) or macrophages differentiated from HoxB8-immortalized progenitor cells (F) were subjected to polysome profiling analysis. Total protein was extracted from individual fractions and subjected to Western blot analysis for the indicated proteins. Puromycin and RNase treatments were performed as described in the Methods section.

Intriguingly, among novel RAPs that were highly enriched in TLR3-stimulated macrophages, we identified multiple cytoplasmic RNA sensors. Two such sensors were OAS3 and OAS1a, members of the 2’,5’-oligoadenylate synthetase (OAS) family that are known to regulate translation in the context of viral infection but have not been previously identified as RAPs. OAS3 and OAS1a detect intracellular viral dsRNA and activate RNase L which, in turn, is thought to nonspecifically degrade ribosomes (rRNAs) and viral RNA^89^. Other identified sensors included RIG-I, LGP-2, and MDA5 which result in type I interferon induction upon ligand recognition^90^; PKR which can induce translational inhibition^91^; and ZBP1 which can induce cell death^92, 93^. It is intriguing to speculate that these findings indicate a mode of on-ribosome sensing of immunostimulatory viral RNAs which may both 1) increase the likelihood of encounters between sensors and ligands and 2) provide a mechanism of selectively degrading ‘infected’ ribosomes via local OAS-mediated activation of RNase L while allowing ‘uninfected’ ribosomes to proceed with translation of host-derived antiviral mRNAs. Indeed, in support of our proteomic results, published work has confirmed the ability of one sensor, RIG-I, to localize to ribosomes via binding to an rRNA expansion segment on the 60S subunit^94^.

In addition to RNA sensors, we identified several other interferon-stimulated genes with enigmatic functions as constituents of ribosome complexes in TLR3-activated macrophages. Among them, Viperin (also known as Rsad2) and CMPK2 were particularly notable as they have both recently been implicated in translational regulation^95, 96^. Mechanistically, these proteins are required for the generation of the CTP metabolite 3’-Deoxy-3’,4’-didehydro-cytidine triphosphate (ddhCTP) which has been proposed to inhibit viral translation via induction of ribosome collisions. The presence of Viperin and CMPK2 in ribosome complexes may, again, suggest a more localized inhibition of translation than previously thought, whereby CMPK2 and Viperin-bound ribosomes may experience collisions while CMPK2 and Viperin-free ribosomes proceed with unimpeded translation.

Finally, given that TLR3-remodeled ribosomes appeared poised to nucleate signaling pathways downstream of viral RNA detection, we asked whether the TLR4 agonist bacterial LPS would remodel ribosome complexes in a different manner for the distinct purpose of promoting an antibacterial response. Strikingly, however, stimulation of BMDMs with LPS for 0, 2, 6, 12, and 24 hours showed an overall remarkably similar effect on ribosome complexes as did activation with poly(I:C) (Figure 5A and B, Supplementary Figure 3B, Supplementary Table 2). Of the 198 proteins enriched 2-fold, 107 were also 2-fold enriched, and an additional 19 were 1.5-fold enriched following poly(I:C) stimulation. Among RAPs ejected from ribosomes upon LPS, we again observed PDCD4 and HMGB proteins. This similarity in ribosome remodeling may be due to the fact that TLR3 and TLR4 signal through a shared downstream adapter protein, TRIF, which coordinates type I interferon induction and may similarly coordinate ribosome complex reorganization (TLR4, but not TLR3, additionally signals through the adapter MyD88). Indeed, many of the novel RAPs we identified were themselves interferon-stimulated genes (ISGs), and the greater number of highly-enriched RAPs identified following poly(I:C) stimulation (395) compared to LPS stimulation (198) may reflect the more robust TRIF-dependent signaling and interferon induction which occurs downstream of TLR3 ligation.

We performed sucrose gradient fractionation followed by western blotting of collected fractions to confirm several hits common to TLR3- and TLR4-stimulated macrophages as *bona fide* RAPs, including USP15, HO-1, OAS3, Viperin, and CMPK2 (Figure 5C-F) As expected, *in vitro* treatment of activated macrophages with the polysome-dissociating drug puromycin or treatment of lysates with RNaseA/T1 resulted in polysome ‘collapse’ and loss of ribosomal subunits from the heavy fractions. Importantly, puromycin and RNase treatments also resulted in loss of the RAPs from heavy fractions together with concomitant enrichment in the monosome peak, confirming their specific association with the core ribosomal machinery.

Lastly, we did identify several intriguing RAPs that, although detected in both TMT-MS datasets, were selectively enriched following LPS treatment. These included the relatively uncharacterized protein Mndal^97^, as well as CARD9 and SON, which have known roles in immune defense^98, 99^. Both SON and CARD9 have been putatively identified as RAPs^88^. We also identified the endoplasmic reticulum (ER)-resident protein Sec61b as selectively enriched following TLR4 stimulation. Ribosomes are known to localize to the outer ER membrane via interaction with Sec61b^10^. Preferential interactions following LPS may suggest relocalization of ribosomes to the ER, or increased synthesis of ER-bound ribosomes, in order to translate the robust quantities of cytokines that are elaborated following this particular challenge.

Thereby, RAPIDASH has enabled the characterization of dynamic changes to ribosomes upon macrophage stimulation and serves as a resource for the identification and characterization of how ribosome remodeling, at the level of RAPs, may drive very rapid changes in control of the translatome.

## Discussion

We have developed a method called RAPIDASH that can enrich ribosomes and their associated proteins. The specificity of RAPIDASH is due in large part to the second step of chromatography with cysteine-charged sulfolink resin. Although the cysteine-charged sulfolink resin preferentially binds to rRNA compared to poly(A) RNA, the mechanism by which this specificity is achieved is not clear and could potentially be due to an affinity for structured RNA or RNA modifications that are present in rRNA.

Compared to Ribo-Flag IP, RAPIDASH can be used on any sample, requires less input material, and has better coverage. We have demonstrated that RAPIDASH can successfully identify RAPs in different cells and tissues, as well as upon stimuli, and we believe RAPIDASH can be used for many other cell types, tissues, organisms, and dynamic conditions, including clinical settings and non-model organisms. RAPIDASH is capable of processing low input samples, enriching on average about 350 ng of RAPs from a single E12.5 limb bud. In addition, while RAPIDASH can generate a decent quality list of candidate RAPs, it is important to validate that these proteins are true hits, for example, by sucrose gradient fractionation of samples treated with EDTA.

Previous work described pervasive translational regulation of cell signaling components that are important for mouse development, but it was unclear how the ribosome itself played a role. Our findings in tissues suggest that RAPs may enable the ribosome to coordinate tissue-specific functions. For example, the helicase function of Dhx30 may regulate the translational specificity of the ribosome in neurons by denaturing secondary structures of select mRNAs; it would be interesting to identify the targets of Dhx30 and how their translational efficiency changes if Dhx30 carries the same mutations as in patients. Our data suggest LLPH may enable the proficient translation of neural developmental genes with long coding sequences, including those that are required for neurite outgrowth, revealing a potential new mechanism for neural translation.

From our relative quantification mass spectrometry data, we have identified various RAPs that could play a tissue specific role in normal mouse development. Elavl2 is capable of binding both ribosome and mRNA; it remains to be seen precisely how this impacts translation of Elavl2 binding targets. Interestingly, we see cytoskeleton-related components such as Vimentin and Map1b being enriched in the forebrain. Other cytoskeletal components have previously been found to be associated with ribosomes, which hints at possible roles in localized translation^100–103^. Map1b has been shown to play a role in microtubule remodeling critical for axonal outgrowth^104^. Possibly, ribosomes are recruited alongside Map1b to distal growth cones, or facilitate central sprouting and peripheral regeneration of neurons^104–106^. Additionally, the identification of Ckb and Ptprz1, a kinase and phosphatase, respectively, as potential RAPs suggests they enable rapid translational responses to cell signals. Finally, we have applied RAPIDASH to stimulated macrophages over time and observed extensive changes in ribosome composition. These data suggest that pathogen detection leads to a translational rewiring to help macrophages overcome infection. Notably, activation of TLR3 and TLR4 lead to overlapping changes in ribosome composition, which may suggest the usage of common mechanisms of translational control despite distinctions in inflammatory contexts.

Thus, RAPIDASH has revealed a dynamic and diverse layer of ribosome composition that is important for tissue-specific functions and responses to acute stimuli. In the future, RAPIDASH can be used to characterize ribosome composition in a variety of samples to characterize how the ribosome acts as a cell signal integrator to regulate gene expression in health and disease.

## Acknowledgements

This work was funded by National Institute of Health (NIH) grants 5R21MH130323 and 5R01HD086634 (to M.B.), R38 HL143581 and K38 HL165490 (to A.G.L.), ad R35CA242986 (to D.R.). D.R. was also supported by the American Cancer Society (Research Professor Award). Mass spectrometry for the LLPH-FLAG IP-MS experiment and for macrophages was provided by the Mass Spectrometry Resource at UCSF (A.L.B., Director) supported by the Dr. Miriam and Sheldon G. Adelson Medical Research Foundation (AMRF) and the UCSF Program for Breakthrough Biomedical Research (PBBR). T.T.S. was supported by a National Science Scholarship (PhD) from the Agency for Science, Technology and Research. V.H. was the Fraternal Order of Eagles Fellow of the Damon Runyon Cancer Research Foundation (DRG 2314-17) and was also supported by the Katharine McCormick Advanced Postdoctoral Fellowship. C.H.K. was supported by a Stanford Dean’s Fellowship and a Canadian Institutes of Health Research Postdoctoral Fellowship. Y.Y. was supported by the New York Stem Cell Foundation Druckenmiller Fellowship (NYSCF-D-F74). Y.C. was supported by a Stanford Dean’s Fellowship. L.F. was supported by an EMBO Long-Term Fellowship. K.F. was supported by the Uehara Memorial Foundation and Human Frontier Science Program Fellowship.

## Author Contributions

T.T.S., V.H. and A.G.L. contributed equally to this work. T.T.S., V.H., K.F., and M.B. developed the RAPIDASH technique. T.T.S., V.H., and Y.C. characterized the cysteine-charged sulfolink resin. T.T.S. and V.H. designed and performed the proteomic characterization and analysis of mESCs and mouse embryonic tissues. C.H.K. performed the characterization of Dhx30. L.F. performed the characterization of LLPH in P493-6 and A549 cells. Y.Y. and M.W. aided the design of the experiments in hiNs. T.T.S. performed the characterization of LLPH in hESCs and hiNs. A.G.L. and D.R. designed the macrophage experiments. A.G.L. performed the macrophage experiments. J.A.O.-P. and A.L.B. helped design and also performed the mass spectrometry and analysis of the LLPH IP-MS data and the macrophage data. T.T.S., V.H., A.G.L., M.B., and D.R. wrote the paper. All authors edited the paper.

## Materials and Methods

### Mice

#### Embryonic tissue-related experiments

Mouse protocols were reviewed and approved by the Stanford Administrative Panel on Laboratory Animal Care (APLAC, protocol #27463). Mice were housed at Stanford University with standard conditions: 12 hour light-dark cycle, ambient temperatures between 68 and 79 °F, humidity between 30 and 70%, free access to chow, acidified water, and filtered air flow. Wild-type FVB/NJ mice were purchased from Jackson Laboratories. For timed pregnancies, one male and one female were housed together, and the female was checked daily for plugs. The day the vaginal plug was observed was considered embryo stage E0.5. At E12.5, the pregnant female was euthanized by CO_2_ inhalation followed by cervical dislocation to collect the embryos.

#### Macrophage-related experiments

Wild-type C57BL/6J mice were purchased from Jackson Laboratories. Animals were housed in a specific-pathogen free environment in the Laboratory Animal Research Center at UCSF. All experiments conformed to ethics and guidelines approved by the UCSF Institutional and Animal Care and Use Committee.

### Cell culture

Wild-type E14 mouse embryonic stem cells (mESCs) were a gift from Barbara Panning’s lab (UCSF). The Rpl36-FLAG mESC and Rps17-FLAG mESC lines were as described previously^10^. All mESC lines were maintained on cell culture dishes coated with 0.1% (w/v) gelatin in sterile milliQ water in cell culture incubators at 5% CO_2_, 37 °C in mESC media containing KnockOut^TM^ DMEM (Dulbecco’s Modified Eagle Media; Thermo Fisher Scientific, 10829-018) with 15% ES quality fetal bovine serum (FBS) (Millipore, ES-009-B), 2 mM EmbryoMax® L-glutamine (Millipore, TMS-002-C), 1x EmbryoMax® Penicillin/Streptomycin (Millipore, TMS-AB2-C), 1x EmbryoMax® MEM, non essential amino acids (Gibco, 11140050), 0.055 mM 2-mercaptoethanol (Gibco, 21985023), and 10^3^ U/ml mouse leukemia inhibitory factor (LIF) protein (Gemini, 400-495). Cells were typically passaged every other day at a 1:6 ratio for maintenance. On days where the cells were not split, the media was changed to fresh mESC media.

For differentiation of primary BMDMs, whole bone marrow from femurs and tibias was plated on polystyrene dishes (five 10-cm dishes per mouse) for six days in complete DMEM (10% heat-inactivated FBS, 10mM HEPES (Gibco), 100 U/mL Penicillin, 100 µg/mL Streptomycin (Gibco), 1 mM Sodium Pyruvate (Gibco), 1x Glutamax (Gibco), and 55 µM 2-mercaptoethanol (Gibco)) supplemented with 10% mCSF medium derived from 3T3-mCSF cells (a kind gift from Hiten Modani’s lab at UCSF). BMDMs were replated overnight on tissue-culture-treated dishes prior to activation with LPS (100 ng/mL, Invivogen) or poly(I:C) (2 µg/mL, Invivogen).

For some experiments, macrophages were generated from HoxB8-immortalized murine progenitor cells (a kind gift from Averil Ma’s lab at UCSF). HoxB8 cells were maintained in an undifferentiated state in complete RPMI (10% heat-inactivated FBS, Glutamax, HEPES, Penicillin/Streptomycin) supplemented with b-estradiol (1 µM) and 5% Flt3L medium derived from B16-FLT3 cells. For macrophage differentiation, HoxB8 progenitor cells were washed twice in complete RPMI without b-estradiol and plated on polystyrene dishes in complete DMEM supplemented with 10% mCSF medium for six days. On day six, macrophages were replated onto tissue-culture treated dishes overnight prior to activation.

### Characterization of cysteine-charged sulfolink resin (related to Figure 1B)

#### mESC ribosome cushion pellet preparation

mESC pellets were lysed in 350 µL cold <u>lysis buffer A</u> (20 mM HEPES (Sigma-Aldrich, H3375-100G), adjusted to pH 7.6 by KOH (Sigma-Aldrich, 221473-500G) (HEPES-KOH), 15 mM magnesium acetate (Mg(OAc)_2_) (Sigma-Aldrich, M5661-250G), 60 mM NH_4_Cl (Sigma-Aldrich, A9434-500G), 1 mM dithiothreitol (DTT; Pierce, A39255), 100 µg/ml cycloheximide (CHX; Sigma-Aldrich, C7698-1G), 1% Triton X-100 (Sigma-Aldrich, T8787), 0.5% sodium deoxycholate (Sigma-Aldrich, D6750), 8% glycerol (Sigma-Aldrich, G9012-100ML), 20 U/ml Turbo DNase (Ambion, AM2238), 200 U/ml SUPERase-In RNase Inhibitor (Ambion, AM2696), 1× Halt™ Protease and Phosphatase Inhibitor (Thermo Scientific, 78443), in nuclease-free water (Invitrogen, 10977015)) at 4 °C for 30 minutes with occasional vertexing every 10 min. Lysates were cleared by sequential centrifugation at 800 ×g for 5 minutes, 800 ×g for 5 minutes, 8000 ×g for 5 minutes and 21,300 ×g for 5 minutes. Cleared supernatant were loaded on to 700 μL 1 M sucrose <u>cushion buffer</u> (20 mM HEPES-KOH, pH 7.6, 15mM Mg(OAc)_2_, 60 mM NH_4_Cl, 1 mM DTT, 100 µg/ml CHX, 200 U/ml SUPERase-In RNase Inhibitor, 1 M sucrose (Fisher Scientific, S5-12)) and centrifuge at 100,000 rpm at 4 °C using Beckman TLA100.3 rotor for 1 hour. Ribosome cushion pellets were gently washed and resuspended in 300 μL <u>binding buffer</u> (20 mM HEPES-KOH, pH 7.6, 15 mM Mg(OAc)_2_, 60 mM NH_4_Cl, 1 mM DTT, 100 µg/ml CHX, 200 U/mL SUPERase In RNase inhibitor, 20 U/mL Turbo DNAse,) at 4 °C in a Thermomixer with 1,000 rpm for 1 hour. Ribosomal RNA concentration was quantified by Qubit RNA HS kit (Life Technologies, Q32852) following manufacturer’s instructions.

#### Isolation of poly(A) RNA

Total RNA was first extracted from mESCs. A frozen cell pellet from a confluent 10-cm plate of mESCs was thawed on ice, resuspended in 1 ml cold TRIzol, and incubated at room temperature for 5 minutes. Then, 200 µL chloroform (Millipore Sigma, 3150) was added, and the tube was manually shaken for 15 seconds. The tube was then incubated at room temperature for 2 minutes. The sample was centrifuged at 12,000 ×g for 15 minutes at 4 °C. Then 510 µL top aqueous layer was pipetted out into a fresh, chilled tube. This was then mixed with 510 µL 70% ethanol. The RNA was purified using a Purelink RNA Mini kit (Thermo Fisher, 12183025) following the manufacturer’s instructions with the following exceptions. The sample was transferred to a PureLink spin cartridge and centrifuged at 12,000 ×g for 15 seconds at room temperature. The flowthrough was discarded, and the cartridge was reinserted into the same collection tube. This centrifugation step was repeated so the entire sample was bound to the cartridge. Washes were performed according to the manufacturer’s protocol. To elute, the cartridge was inserted into a DNA lo-bind tube, and 50 µL RNase-free water was added to the center of the cartridge. The sample was incubated at room temperature for 1 minute. Finally, the sample was centrifuged at 12,000 ×g for 1 minute at room temperature. The resulting eluate was stored at −80 °C.

Poly(A) RNA was enriched using NEBNext® Poly(A) mRNA Magnetic Isolation Module (New England Biolabs (NEB), E7490L) according to the manufacturer’s instructions with some modifications. Each sample that was destined to be incubated with cysteine-charged resin derived from 8 poly(A) reactions with 5 µg RNA each. All steps except for the elution were performed as described in the manufacturer’s protocol up until the final elution. For the final elution, the beads from 8 reactions were combined and resuspended in 60 µL RNase-free water by pipetting six times. The sample was incubated at 80 °C for 2 minutes and then cooled to 25 °C to elute the poly(A) RNA. The tube was then placed on a magnetic rack for about 2 minutes, or until the solution was clear. Then, 57 µL of supernatant was collected and stored at −80 °C.

#### Chromatography with cysteine-charged resin

To perform the cysteine-charged sulfolink characterization, sucrose cushion pellet and poly(A) samples were thawed on ice, and the RNA concentrations were measured using a Qubit RNA high sensitivity (HS) assay kit (Thermo Fisher, Q32852). Samples were diluted to RNA concentrations in the range of 4.53 - 7.07 ng/µL using RNase-free water and then kept on ice while the resin was prepared.

First, 500 µL Sulfolink coupling resin (Thermo Fisher, 20402) was added to a falcon tube and centrifuged at 850 ×g for 1 minute at room temperature, and the supernatant was decanted. The resin was then resuspended in 500 µL <u>coupling buffer</u> (50 mM Tris-HCl, pH 8.5 (Fisher Scientific, BP153-500), 5 mM ethylenediaminetetraacetic acid (EDTA; Thermo Fisher, AM9262)), and then centrifuged at 850 ×g for 1 minute at room temperature. The supernatant was decanted, and the resin was washed twice more with 500 µL coupling buffer as previously described. After the last wash was decanted, the resin was resuspended in 500 µL 50 mM L-cysteine (Sigma, 168149-25G) in coupling buffer and left on a rocker at room temperature for at least one hour. Then, the resin was centrifuged at 850 ×g for 1 minute at room temperature. The supernatant was decanted, and then the resin was resuspended in 500 µL coupling buffer and centrifuged at 850 ×g for 1 minute at room temperature. The supernatant was decanted, and this washing step was repeated twice more. The resin was then washed three times with 1 mL <u>priming buffer</u> (20 mM HEPES-KOH, pH 7.6, 15 mM Mg(OAc)_2_, 60 mM NH_4_Cl, 1 mM DTT, and 100 µg/mL CHX) by resuspending the resin in the priming buffer, centrifuging at 850 ×g for 1 minute, and decanting the supernatant. The wash with priming buffer was repeated twice more, and then the resin was resuspended in 500 µL priming buffer. For each sample, 12.5 µL resin slurry was added to a micro-spin column (Thermo fisher, PI89879), which was placed in a DNA lo-bind tube. The caps were removed. The micro-spin columns were centrifuged at 1,000 ×g for 1 minute at room temperature to remove the priming buffer from the resin, and then the bottom caps were put on again.

For each sucrose cushion pellet or poly(A) RNA sample, 42 - 48 µL was added to the resin for a total of 190 - 339 ng of RNA. The top caps were screwed on, and the columns were briefly vortexed to mix. The capped columns were put in DNA lo-bind tubes and put on ice for 15 minutes under foil. Then, the caps were removed, the columns were moved to fresh DNA lo-bind tubes, and the samples were centrifuged at 1,000 ×g for 1 minute at room temperature. The bottom caps were replaced, the flowthroughs were re-applied to the resin, and the top caps were screwed on. The columns were again briefly vortexed to mix, and then they were returned to the DNA lo-bind tubes and left on ice for 15 minutes under foil. Then, the caps were removed, and the samples were centrifuged at 1,000 ×g for 1 minute at room temperature. The flowthroughs were kept on ice, and the bottom caps were put on to the columns, which were moved to fresh DNA lo-bind tubes. The resin was then washed by adding 12.5 µL priming buffer to each column, capping the columns, vortexing the columns briefly to mix, uncapping the columns and returning them to their DNA lo-bind tubes, and centrifuging the samples at 1,000 ×g for 1 minute at room temperature. This washing step was repeated three more times for a total of four washes. The bottom caps of the columns were put on, and the columns were moved to fresh DNA-lo bind tubes. To elute, 12.5 µL <u>elution buffer</u> (20 mM HEPES-KOH, pH 7.6, 15 mM Mg(OAc)_2_, 500 mM KCl (Invitrogen, AM9010), 1 mM DTT, and 100 µg/mL CHX) was added to each column. The top caps were screwed on, and the columns were vortexed briefly to mix and then returned to their DNA lo-bind tubes. The samples were then rested on ice for 2 minutes under foil. Finally, the caps were removed, and the eluates were collected by centrifuging at 1,000 ×g for 1 minute at room temperature. The eluates were kept on ice.

After collecting eluate, 50 µL Trizol (Invitrogen, 15596026) was added to the resin and incubated at room temperature for 5 min. Trizol was collected by centrifuge at 1,000 ×g for 1 min at room temperature. RNA was extracted from each resin using RNA Clean and Concentrator-5 kit (Zymo, R1016). The RNA was eluted in 10 µL RNase-free water, and RNA concentrations were measured by Qubit RNA HS Kit.

### RAPIDASH for E14 mouse embryonic stem cells (related to Figure 1)

#### Harvest and cytoplasmic lysis

Approximately 15 x 10^6^ cells each of FLAG-tagged S17 and FLAG-tagged L36 mESCs as characterized previously^10^ were seeded on to 15-cm plates (1:4 ratio from ∼80% confluent plate). After one day, the media was replaced with 18 mL fresh mESC media. One hour post-media change, 2 mL 1 mg/mL CHX in mESC media was added to each 15-cm plate for a final concentration of 100 μg/mL CHX, and the plates were returned to the cell culture incubator for 3 minutes. The cells in each 15-cm plate were rinsed twice with pre-warmed 10 mL 100 μg/mL CHX in Dulbecco’s Phosphate-Buffered Saline (DPBS; Gibco, 14190-250) and were dissociated with 4 mL 100 μg/mL CHX, 0.05% trypsin-EDTA solution in DPBS. The plates were returned to the incubator briefly, after which the trypsin solution was quenched by adding 4 mL 100 μg/mL CHX in cold mESC media. The cells were pelleted in a tabletop centrifuge at 200 ×g at 4 °C for 3 minutes, and the supernatant was aspirated out. Each pellet was washed once with 2 mL cold 100 μg/mL CHX in DPBS, transferred to a 2 mL tube, and then pelleted again in a tabletop centrifuge at 200 ×g at 4 °C for 3 minutes. The supernatant was aspirated out, and each pellet was flash frozen in liquid nitrogen and stored at −80 °C. Each pellet was lysed in 400 μL cold <u>lysis</u> <u>buffer A</u>. Cells were then lysed by vortexing at high speed for 30 seconds and putting them back on ice for 30 seconds. This was repeated another two times for a total of 3 minutes. The samples were then incubated on ice for 30 minutes and vortexed briefly for 10 seconds every 10 minutes. The cytoplasmic fraction was enriched by centrifuging the lysates twice at 800 ×g for 5 minutes at 4 °C, then once at 8,000 ×g for 5 minutes at 4 °C, and finally once at 20,817 ×g for 5 minutes at 4 °C, with the supernatants being moved to fresh chilled tubes after each centrifugation.

#### Sucrose cushion ultracentrifugation

High density complexes were enriched by sucrose cushion ultracentrifugation. To do this, 300 μL of cytoplasmic lysate from each sample was layered on to 700 μL of <u>sucrose cushion buffer</u> in a polycarbonate centrifuge tube (Beckman Coulter, 343778) and centrifuged using a TLA120.2 rotor (Beckman Coulter, 357656) at 100,000 rpm for 1 hr at 4 °C.

#### Preparation of L-cysteine charged sulfhydryl resin

The amount of L-cysteine charged sulfhydryl resin required for each experiment was prepared while the samples were undergoing sucrose cushion ultracentrifugation. Per sample, 1 mL of SulfoLink™ coupling resin slurry (Thermo Scientific, 20402) was pipetted into a polypropylene tube (Falcon, 14-949-11B), which was used in order to minimize resin loss during decanting. The slurry was initially centrifuged at 800 ×g for 1 minute at 4 °C to pellet the resin, and the storage solution was decanted. The resin was then washed three times by adding a volume of cold coupling buffer equal to twice the bed volume, resuspending the resin by inverting gently, centrifuging at 800 ×g for 1 minute at 4 °C, and decanting the supernatant after each centrifugation. Afterwards, a volume of 50 mM L-cysteine in coupling buffer equal to twice the bed volume was added to the resin. The tube was wrapped in aluminum foil and then rocked for 1 hour on a platform rocker at room temperature. The resin was then centrifuged at 800 ×g for 1 minute at 4 °C, and the supernatant was decanted to collect the L-cysteine charged sulfhydryl resin. The charged resin was washed three times with cold <u>priming buffer</u>. For each wash, cold priming buffer with four times the bed volume was added to the resin, which was resuspended by gently inverting the tube and then collected by centrifuging at 800 ×g for 1 minute at 4 °C and decanting the supernatant. After the final wash, the charged resin was resuspended in a volume of the cold priming buffer equal to twice the bed volume, 1.5 mL charged resin slurry was aliquoted into one 5 mL spin column (Pierce, PI89897) for each sample. Charged resin can be stored overnight at 4 °C.

#### Affinity enrichment with L-cysteine charged sulfolink resin

After the sucrose cushion ultracentrifugation step, the supernatant was removed by pipetting with a P1000 micropipette without disturbing the pellet, which contains high density protein complexes. Each pellet was dislodged with a pipette tip and resuspended in 1 mL of cold <u>binding buffer</u> by pipetting for 30 seconds. The tube was then sealed with parafilm wrap and shaken on a thermomixer at 1,000 rpm for 30 minutes at 4 °C to further resuspend the protein. Immediately prior to incubation, the twist-off bottoms of the spin columns containing the charged resin were snapped off, and the columns were placed in 15 mL falcon tubes. The columns were then centrifuged at 1,000 ×g for 1 minute at 4 °C to remove the priming buffer and subsequently capped with the provided bottom closures. The priming buffer was discarded. Then, for each sample, half of the resuspended protein pellet was transferred into a charged resin-containing spin column. The other half of each sample was retained to analyze the sucrose cushion ultracentrifugation step. The spin columns were flicked several times to mix and then incubated on ice for 15 minutes under foil. The bottom closures were then removed, and the flowthroughs were collected by centrifuging at 1,000 ×g for 1 minute at 4 °C. The spin columns were capped with the bottom closures and replaced into the 15 mL falcon tubes, and the flowthroughs were added back to their respective resins. The resin was resuspended by flicking the spin columns and then incubated for another 15 minutes on ice under foil. The bottom closures were removed, the centrifugation step was repeated, and the flowthroughs were set aside for other analyses. The resin in each spin column was then washed with a volume of cold priming buffer equal to twice the bed volume by capping the spin column with a bottom closure and inverting by hand several times. The bottom closures were then removed, and the columns were centrifuged again at 1,000 ×g for 1 minute at 4 °C to discard the wash buffer. This washing step was repeated three more times. After the final wash, the bottom closures were reattached to the columns. Each sample was then eluted by adding a volume of cold <u>elution buffer</u> equal to half the bed volume, resuspending the resin by flicking the tubes gently, incubating on ice for 2 minutes under foil, and centrifuging the samples at 1,000 ×g for 1 minute at 4 °C. The elution step was repeated three more times with volumes of fresh elution buffer equal to half the bed volume each time. All four elutions for each sample were pooled together.

### RAPIDASH for E12.5 tissues (related to Figure 4B)

Limbs, forebrain, and liver tissues were microdissected from E12.5 FVB/NJ mouse embryos. Uterus was washed in PBS twice to remove the blood. Each embryo is transferred into a plate containing filming media one by one, where microdissection was performed to take out the tissues. Filming media (10% FBS (Sigma-Aldrich, TMS-013-B) in DMEM/F-12 with HEPES without phenol red (Gibco, 11039021)) was used during microdissection, and the tissues were collected with P1000 pipette. Excess media was removed by pipette without centrifugation, and the sample was then flash frozen in liquid nitrogen. Each biological replicate consisted of tissues pooled from the same litter (9-12 embryos).

For lysis, 100 μL of cold <u>lysis buffer A</u> was initially added into each sample. The tissues were rapidly ground by hand for 30 seconds using a disposable microcentrifuge pestle. Then, an additional 300 μL of cold <u>lysis buffer A</u> was added to each sample. RAPIDASH was then performed as described above starting from the vortexing step during lysis (RAPIDASH for E14 mouse embryonic stem cells), with the following modifications: after the sucrose cushion ultracentrifugation step, each protein pellet was resuspended in 500 μL cold binding buffer instead of 1 mL, and the entire volume was subjected to affinity enrichment with L-cysteine charged sulfhydryl resin.

### Mass spectrometry of unlabeled tryptic digests (related to Figures 1G, 1H, and 2A)

#### Protein precipitation

Proteins were precipitated from mESC samples by adding one volume equivalent of −20 °C Precipitation Agent from the ProteoExtract® Protein Precipitation Kit (Millipore, 539180) to each sample. For RAPIDASH samples from E12.5 mouse tissues, four volume equivalents of Precipitation Agent was mixed with each sample instead. All samples were then placed at −20 °C for at least 16 hours. Precipitated protein samples were then pelleted by centrifugation at 10,000 ×g for 10 minutes at room temperature. The supernatant was then removed from each sample by pipetting and discarded. Each protein pellet was then washed with 1 mL of prepared Wash Solution from ProteoExtract® Protein Precipitation Kit that was chilled at −20 °C. The samples were vortexed briefly and centrifuged at 10,000 ×g for 2 minutes at room temperature. The supernatant was removed from each sample, and the wash and centrifugation step was repeated one more time. The pellets were then dried. by leaving the tubes open to allow the remaining wash buffer to evaporate.

#### Trypsin/Lys-C digestion

Each protein sample was solubilized by pipetting with a low retention pipette tip (Fisher Scientific, 02-717-135) in 50 µL denaturing buffer (50 mM NH_4_HCO_3_ (Sigma Aldrich, 09830-500G), 6 M urea (Sigma Aldrich, U1250-1KG), in high performance liquid chromatography (HPLC) water (Fisher Scientific,W5-4)). The protein concentration for each sample was then quantified using a Bradford assay (Bio-Rad, 5000006) with technical duplicates following the manufacturer’s protocol. For each mESC RAPIDASH sample (Figures 1G-H), 80 μg protein was denatured and reduced by adding 0.5 µL 500 mM DTT to each sample for a final concentration of 5 mM and incubating in a thermomixer at 37 °C for 1 hour at 500 rpm. The samples were alkylated by adding 1 µL 500 mM iodoacetamide (Pierce, A39271) to each 50 µL sample for a final concentration of 15 mM and incubating them in the dark at room temperature for 30 minutes. A 0.2 µg/µL Trypsin/Lys-C stock solution in Resuspension Buffer (Promega, V5073) was subsequently added to each sample at a 1:50 trypsin/Lys-C:sample ratio by mass to digest the protein into tryptic peptides. The samples were mixed and then shaken at 500 rpm for 4 hours in a thermocycler at 37 °C for the initial digestion by Lys-C. Afterwards, each sample was diluted sixfold by adding 250 µL 50 mM NH_4_HCO_3_ to reduce the urea concentration to 1 M to allow trypsin to refold. The samples were then incubated for another 12 hours in a 37 °C water bath to complete the digestion. Trypsinization was then quenched by adding 3 µL 50% v/v heptafluorobutyric acid (HFBA; Sigma-Aldrich, 52411-5ML-F) in HPLC water to each sample. The samples were then either stored at −80 °C or immediately desalted.

#### Peptide desalting

Each sample was desalted using an OMIX C18 pipette tip (Agilent technologies, A57003100) attached to a P200 pipette set to 150 µL. Each OMIX C18 pipette tip was conditioned by pipetting 50% (v/v) acetonitrile up and down three times. No air was allowed to pass through the sorbent after it was conditioned. The OMIX C18 pipette tip was then equilibrated with 1% (v/v) HFBA by pipetting up and down three times. Acidified peptides were bound to the OMIX C18 sorbent by pipetting up and down for seven times carefully. After the peptides were bound, each tip was rinsed with 0.1% (v/v) HFBA by pipetting up and down three times. To elute the peptides, 7 µL 0.1% (v/v) formic acid (Fisher Scientific, A117-50) in 50% (v/v) acetonitrile by carefully pipetting up and down five times. A second elution was performed by pipetting up and down with 7 µL 0.1% (v/v) formic acid in 75% (v/v) acetonitrile five times. The two elutions were then pooled together and dried completely using a speed-vac (∼45 minutes). The dried peptide samples were stored at −80 °C. The peptides were resuspended in 15 µL 0.1% formic acid for analysis by mass spectrometry.

#### Liquid chromatography tandem mass spectrometry

Mass spectrometry was performed on an Acquity UPLC (ultra performance liquid chromatography M-class system (Waters) coupled online to an Orbitrap Elite mass spectrometer (Thermo Fisher Scientific). Peptides were separated by reversed-phase chromatography using a Self-Pack PicoFrit column (New Objective, PF360-75-15-N-5) with a 360 µm outer diameter, 75 µm inner diameter, and a tip size of 15 µm packed to approximately 22 cm with HALO Peptide ES-C18 Bulk Packing 2.7 µm beads (MAC-MOD Analytical, 942120202). The UPLC solvents A and B were 0.1% (v/v) formic acid and 0.1% formic acid (v/v) in/100% acetonitrile, respectively. For each sample, 2 µL was loaded at 1% B at 0.3 µL/minute for 20 minutes. Peptides were then separated at 0.3 µL/minute over a linear gradient from 1% B to 40% B for 90 minutes, followed by a linear gradient from 40% B to 100% B for 20 minutes, followed by a constant flow at 100% B for 10 minutes. Then, there was a linear ramp back down to 1% B over 5 minutes and then constant flow at 1% B for 5 minutes.

The Orbitrap Elite (Thermo Fisher Scientific) was operated in a data-dependent mode using Xcalibur v3.0 (Thermo Fisher Scientific). Collision-induced dissociation (CID) MS/MS scans were recorded after each MS1 scan (R = 120,000) for the top 15 most abundant precursor ions in the Orbitrap. The isolation width was 1.8 m/z, the normalized collision energy was 35%, and the activation time was 10 milliseconds. The dynamic exclusion parameters were: a repeat count 1 with a 45 second repeat duration, an exclusion list size of 500, an exclusion duration of 80 seconds, and a +/- 10 ppm exclusion mass width. Only charge states of 2 or 3 were not rejected; others, including unassigned charges, were rejected.

### Tandem mass tag (TMT) mass spectrometry (related to Figures 1D and 4B)

For the comparison of the original sulfhydryl-charged resin chromatographic purification of eukaryotic ribosomes and the RAPIDASH protocol, mammalian ribosomes were purified from one 70% confluent 10-cm plate of mESCs following the previously published protocol^10^ or using the “RAPIDASH for E14 mouse embryonic stem cells protocol.”

Proteins were precipitated from mESC sucrose cushion pellet, mESC RAPIDASH, or E12.5 mouse tissues RAPIDASH samples as described above (Mass spectrometry from unlabeled tryptic digests).

The protein samples were then digested into peptides as described in (Mass spectrometry from unlabeled tryptic digests - Trypsin/Lys-C digestion). To prepare samples for TMT, equal amounts of protein by mass were aliquoted into separate tubes according to Table 1.

#### TMT labeling

TMTsixplex™ Isobaric Label Reagents (Thermo Scientific, 90066) were freshly prepared for each experiment by resuspending the entire 0.8 mg of labeling reagent in the tube with 100 µL 100% ethanol (Gold Shield, 412804). For every 1 µg of peptide, 0.5 µL 8 µg/µL of freshly resuspended TMT label reagent was used. To do this, the peptides were first resuspended in 20 mM HEPES, pH 8.0 (Fisher Scientific, AAJ63578AK). The volume of HEPES buffer used was three times the volume of the TMT label that would be added. TMT label reagents were then added into the resuspended peptides, and the samples were incubated for 1 hour at room temperature in the dark. The labeling reaction was subsequently quenched by adding a volume of 5% (v/v) hydroxylamine (Sigma Aldrich, 467804-10ML) equal to 1/20 of the sample volume at room temperature for 15 minutes. The samples for a single injection for relative quantification with different TMT labels were then mixed together according to Table 1. The combined sample was then acidified by adding 50% HFBA to a final concentration of ∼0.5% (v/v) and 100% formic acid to a final concentration of 5% (v/v). Then, the combined samples were desalted using OMIX C18 pipette tips and dried with a speed-vac as described earlier. Peptides were resuspended in 8 µL 0.1% (v/v) trifluoroacetic acid (Fluka, 40967) in 5% (v/v) acetonitrile and subjected to ultra performance liquid chromatography (UPLC) tandem mass spectrometry (MS/MS) analysis using an Acquity UPLC M-class system (Waters) coupled online to an Orbitrap Elite mass spectrometer (Thermo Fisher Scientific).

#### Liquid chromatography tandem mass spectrometry for TMT

Peptides were separated by reversed-phase chromatography using the same column as in “Mass spectrometry from unlabeled tryptic digests - Liquid chromatography tandem mass spectrometry”. UPLC solvent A was 0.1% (v/v) formic acid, and UPLC solvent B was 0.1% (v/v) formic acid/100% acetonitrile, respectively. For each sample, 3 µL was injected and loaded for 30 minutes at 1% B at a flow rate of 0.3 µL/minute. Peptides were separated at the same flow rate over a linear gradient of 5% B to 40% B for 180 minutes, followed by a linear ramp to 100% B for 10 minutes, followed by constant flow at 100% B for 10 minutes. Finally, there was a ramp down to 1% B over 1 minute, where it was held for 9 minutes at constant flow.

The Orbitrap Elite (Thermo Fisher Scientific) was operated in a data-dependent mode using Xcalibur v3.0 (Thermo Fisher Scientific). Higher energy collisional dissociation (HCD) MS/MS scans (R = 15,000) were recorded after each MS1 scan (R = 60,000) for the top 15 most abundant precursor ions in the Orbitrap. The following HCD parameters were used: an isolation width of 1.6 m/z, a normalized collision energy of 40%, and an activation time of 0.1 milliseconds. Dynamic exclusion parameters were set as follows: a repeat count 1 with a 30 second repeat duration, an exclusion list size of 500, an exclusion duration of 60 seconds, and a +/- 10 ppm exclusion mass width. Charge state rejection was enabled for charge states that were less than 2 or unassigned.

### Mass spectrometry data analysis (related to Figures 1D, 1G, 1H, 2A, 4B)

The raw spectra were analyzed using MaxQuant (v1.6.5.0) against *Mus musculus* SwissProt reviewed proteome database downloaded on April 28, 2020 with the following parameters: a maximum of 2 missed trypsin enzyme cleavage sites, first search mass tolerance of 20 ppm, main search mass tolerance of 4.5 ppm, and MS/MS match tolerance of 20 ppm. Deamidation (NQ), oxidation (M), and N-terminal acetylation were searched as variable modifications, and carbamidomethyl (C) was searched as a fixed modification. For identification, a minimum of one razor + unique peptide was required, and the result was filtered with 1% false discovery rate (FDR) at the peptide and protein levels. For TMT quantification, a minimum of two razor + unique peptides present was required instead.

For data related to Figure 1G (Supplementary Table 3) and 2A (Supplementary Table 4), proteins that were identified only by site, contaminants, and reverse hits were filtered out. Proteins that were identified in three out of six mESC samples (Figure 1G) or two out of three forebrain samples (Figure 2A) were used for subsequent analysis. Gene ontology molecular function (GOMF) terms were obtained by first converting SwissProt IDs into Entrez Gene IDs. The Entrez Gene IDs were then analyzed by Manteia to obtain gene ontology terms that are significant. These were filtered so that level 4 terms were the most broad term level that was utilized (e.g. for terms that had more than one level associated, those that were level 1, 2, or 3 in any branch were filtered out).

For the TMT mass spectrometry data, contaminants, reverse hits, and peptides only identified by site were filtered out, and reporter intensity for each protein was Log2 transformed and median-normalized. For the data related to Figure 1D and Supplementary Figure 1B (Supplementary Table 2), only proteins that were detected in all three replicates were selected. Subsequently, the proteins were categorized to separate core RPs from other proteins. The significance of enriched protein groups in each replicate was evaluated with Welch’s t-test. For the data related to Figure 4B (Supplementary Table 6), proteins that were detected in three out of four replicates were selected, the missing values were imputed by random numbers from a normal distribution, and the reporter intensity values between each pair of tissues were analyzed with a paired student *t*-test using Perseus (v1.6.5.0) Proteins with a fold change ≥ 2 and a permutation-based FDR < 10% were deemed to be significantly enriched in one E12.5 mouse embryo tissue over another.

### Sucrose Gradient Fractionation for RAPIDASH characterization (related to Figure 1C and Supplementary Figure 1A)

Three 15-cm plates of wild-type mESCs were grown and harvested as described previously (RAPIDASH for E14 mouse embryonic stem cells - Harvest and cytoplasmic lysis). All three cell pellets were flash frozen in liquid nitrogen. One cell pellet was lysed and subjected to sucrose cushion ultracentrifugation (RAPIDASH for E14 mouse embryonic stem cells - Sucrose cushion ultracentrifugation), and the resulting sucrose cushion pellet was resuspended in binding buffer and set aside on ice. The second cell pellet was lysed and subjected to the complete RAPIDASH protocol described above (RAPIDASH for E14 mouse embryonic stem cells). The remaining cytoplasmic lysate from these two samples was combined and analyzed by sucrose gradient fractionation as an input control. The last cell pellet was resuspended in 400 μL binding buffer. The cells were lysed by bead milling with a TissueLyser II (Qiagen, 85300) using a 5 mm stainless steel bead (Qiagen, 69989) for 30 seconds at 25 Hz. The cytoplasmic fraction was then isolated by serial centrifugation, and subjected to affinity enrichment with L-cysteine charged sulfhydryl resin as described above (RAPIDASH for E14 mouse embryonic stem cells - Affinity enrichment with L-cysteine charged sulfhydryl resin).

RNA concentrations were measured using a NanoDrop™ 2000c Spectrophotometer to normalize the amount of RNA loaded for each sample. Linear sucrose gradients (10-45% sucrose (w/v), 20 mM Tris-HCl pH 7.5 (Invitrogen, AM9010), 15 mM MgCl_2_ (Invitrogen, AM9010), 150 mM NaCl (Invitrogen, AM9010), 1 mM DTT, 100 μg/mL CHX, in nuclease-free water) were made in 14×89 mm open-top thinwall polypropylene centrifuge tubes (Beckman Coulter, 331372) using a gradient maker (Biocomp, 108). Before layering the samples onto the gradient, 200 μL sucrose gradient solution was removed from each centrifuge tube. For each experimental sample (sulfhydryl-charged resin sample and RAPIDASH sample), 60 μg RNA in 200 μL of their respective buffers was loaded. For a cytoplasmic lysate sample, 180 µg of RNA was used instead. The samples were centrifuged using a SW 41 Ti swinging bucket rotor (Beckman Coulter, 331336) at 40,000 rpm for 2.5 hours at 4 °C.

Fractions were collected for 30 seconds each using the Density Gradient Fraction System with a flow rate of 1.5 mL/minute (Brandel, BR-188l). Proteins were precipitated from fractionated samples as described above, except 600 μL −20 °C Precipitation Agent was added to each fraction (Mass spectrometry from unlabeled tryptic digests - Protein precipitation)). After the protein pellets were dried, they were immediately prepared for Western Blotting.

### Sucrose Gradient Fractionation for mouse E12.5 tissues (related to Figure 2C)

Each sample consisted of tissues pooled from the same litter prepared as described above (RAPIDASH for E12.5 tissues). However, cold lysis buffer B (20 mM Tris-HCl pH 7.5, 15 mM MgCl_2_, 150 mM NaCl, 1 mM DTT, 100 μg/mL CHX, 1% Triton X-100, 0.5% sodium deoxycholate, 8% glycerol, 0.02 U/μL TURBO DNase, 0.2 U/μL SUPERase In^TM^ RNase inhibitor, 1x Halt^TM^ protease and phosphatase inhibitor in nuclease-free water) was added to each sample instead of cold lysis buffer A. Lysis was performed as described above (RAPIDASH for E12.5 tissues), and the cytoplasmic fraction was enriched by serial centrifugation as described previously (RAPIDASH for E14 mouse embryonic stem cells).

Linear sucrose gradients (Sucrose Gradient Fractionation for RAPIDASH characterization) were made in 11×60 mm open-top polyallomer centrifuge tubes (Seton Scientific, 5010). For EDTA treated samples, 50 mM EDTA was included in the sucrose gradient solution. Before layering the cytoplasmic lysates onto the gradient, 130 μL sucrose gradient solution was removed from each centrifuge tube. Typically, 125 μL cytoplasmic lysate containing 80-120 μg RNA was used for each sucrose gradient fractionation experiment. The tubes were centrifuged using a SW 60 Ti swinging bucket rotor (Beckman Coulter, 335649) at 35,000 rpm for 2.5 hours at 4 °C.

Fractions were collected for 28 seconds each using the Density Gradient Fraction System with a flow rate of 0.75 mL/minute (Brandel, BR-188l). Proteins were precipitated and prepared as mentioned above (Mass spectrometry from unlabeled tryptic digests - Protein precipitation). After the protein pellets were dried, they were immediately prepared for western blotting.

### Western blotting for mESCs, human cells, and mouse embryonic tissues (related to Figures 1-4)

Protein samples were resuspended in 1X sodium dodecyl sulfate (SDS) Laemmli buffer (Fisher Scientific, 50-196-784). Protein samples were then denatured in a thermocycler for 10 minutes at 95 °C. Sucrose gradient fractionation samples were separated on 4–20% Criterion™ TGX™ Gels (Bio-Rad, 5671095), and others were separated on 4–20% Mini-PROTEAN® TGX™ Precast Protein Gels (Bio-Rad, 456-1096). The gels were run with 1x Tris/Glycine/SDS buffer (Bio-Rad, 161-0772) until the bromophenol blue dye front reached the bottom of the gel. Unless otherwise stated, semi-dry transfer was performed using a Trans-Blot® Turbo™ RTA Midi (Bio-Rad, 170-4273) or Mini (Bio-Rad, 170-4272) PVDF Transfer Kit on a Trans-Blot® Turbo™ Transfer System (Bio-Rad, 1704150). Each transferred membrane was rinsed once with PBS+0.1% (v/v) TWEEN® 20 (Sigma Aldrich, P1629-100ML) (PBST) and then blocked in 5% (w/v) bovine serum albumin (BSA; Sigma, A9647-100G) in PBST for 30 minutes in room temperature or overnight at 4 °C. The membranes were rocked in primary antibody solutions in 5% (w/v) BSA, 0.02% (w/v) sodium azide (Sigma Aldrich, S2002-25G) in PBST overnight at 4 °C. After the primary antibody incubation, the membrane was then washed four times with PBST for 5 minutes each time. HRP-conjugated anti-mouse (Cytiva, NA931-1ML) and anti-rabbit (Cytiva, NA934-1ML) secondary antibodies were diluted at 1:5000 in 5% milk in PBST. HRP-conjugated anti-goat antibody (R&D Systems, HAF019) was used at a 1:1000 dilution in 5% milk in PBST. The membranes were rocked in secondary antibody solutions for 1 hour at room temperature. The membrane was then washed again four times with PBST for 5 minutes each time. The membrane was then developed with Clarity Western ECL Substrate (Bio-Rad, 170-5061) or SuperSignal™ West Femto Maximum Sensitivity Substrate (Thermo Fisher Scientific, 34095) for at least 5 minutes before imaging it with a ChemiDoc XRS+ System (Bio-Rad, 1708265). The primary antibodies used in this paper are: Nup62 (Proteintech, 13916-1-AP); Atp5a1 (Proteintech, 14676-1-AP); Tom20 (Proteintech, 11802-1-AP); Rpl4 (Proteintech, 11302-1-AP); Rpl10a (Abcam, ab174318); Rps5 (Abcam, ab58345); Rps26 (Proteintech, 14909-1-AP); Rps27 (Fisher Scientific, PIPA518092); Metap1 (R&D Systems, MAB3537-SP); Ufl1 (Bethyl Laboratories, A303-456A); Upf1 (Proteintech, 23379-1-AP); Ddx1 (Bethyl Laboratories, A300-520A); Nsun2 (Proteintech, 20854-1-AP); Dhx30 (Abcam, ab85687); Rpl29 (Abcam, ab88514); Pabp1 (Cell Signaling Technology, 4992); Rps19 (Abcam, ab181365); β-Actin (8H10D10) (Cell Signaling Technology, 3700S); Rps6 (Cell Signaling Technology, 2217); V5 Tag (Thermo Fisher, R960-25); GAPDH (D16H11) (Cell Signaling Technology, CS5174S); Llph (Invitrogen, PA5-66207); Rps20 (Proteintech, 15692-1-AP); and Elavl2 (Proteintech, 14008-1-AP).

### siRNA-mediated knockdown and OP-puromycin labeling (related to Figure 2E)

Before transfection, 6-well plates were coated overnight with 0.1% Gelatin prior to use. Cultured mESCs were trypsinized, washed, and resuspended in OptiMEM at a density of 5.0 x 10^5^ cells/mL. Cells were then transfected with 25 nM of non-targeting control siRNA #2 (siFluc; Dharmacon, D-001210-02-05) or siRNA target against mouse *Dhx30* (Dharmacon) using Dharmafect Reagent I (Dharmacon, T-2001) as per manufacturer’s protocol. Cells (1 mL) were then plated into each well of a 6-well plate and incubated for 24 h at 37 C in antibiotic-free media. Following incubation, global protein synthesis was measured using O-propargyl-puromycin (OPP). Briefly, cells were labeled with 20 µM of OPP in mESC media for 30 min at 37C.

Following metabolic labeling, cells were harvested and washed twice with 1xPBS. Subsequent cell pellets were resuspended in Zombie Violet Live-Dead Stain (1:500 in PBS; BioLegend, 423113) and incubated for 15 min in the dark. Cells were then washed with Cell Staining Buffer (0.1% NaN3, 2% FBS in HBSS) before being fixed in 1% PFA for 15 min on ice. Cells were then permeabilized in Perm Buffer (0.1% Saponin, 0.1% NaN_3_, 3% FBS in PBS) for 1 hour on ice. Cells were next washed twice with Cell Staining Buffer (without 0.1% NaN_3_), labeled with an Alexa Fluor 555 Picolyl Azide dye (Thermo Fisher, C10642) and incubated for 30 min at room temperature in the dark. Labeled cells were washed and resuspended in Cell Staining Buffer before being analyzed on a LSRII flow cytometer (BD Biosciences) using software packages CellQuest and FlowJo v10.

### Plasmid Transfections (related to Figures 2G and 2H)

Prior to use, 6-well plates were coated overnight with 0.1% Gelatin. For transfections, 250 µL of OptiMEM was mixed with 4 µg of plasmid in one tube and separately 250 µL of OptiMEM was mixed with 10µL of Lipofectamine 2000. Both tubes were incubated for 5 min at room temperature. Following incubation, both tubes were mixed together and incubated for 15 min at room temperature. Cultured mESCs were trypsinized, washed, and resuspended in OptiMEM at a density of 1.0 x 10^6^ cells/mL. After incubation of the Lipofectamine:DNA mixture, 1 mL of cells was added to each tube, mixed gently with a pipet, and immediately transferred to a 6-well plate. After incubation for 4 hours, the media was replaced with complete mESC media. Plates were then incubated at 37C for an additional 24 hours before downstream experiments.

### H1-hESC cell culture and CRISPR (related to Figure 3)

H1-hESCs were cultured in mTeSR1 media (StemCell Technologies, 85850) on plastic dishes coated with Geltrex (Gibco, A1413302). H7-hESCs were passaged 1:10 using Accutase (Gibco, A1110501) and cultured overnight in mTeSR1 supplemented with 2 µM thiazovivin (Tocris, 3845) to promote cell survival.

To generate homozygous Llph knockout in H1 hESCs, guide RNAs (AAGCCTGCTGGCCTCCTTTG, GAGAATACTTTTAAGCCTGC, CTCTTGGCAATGTTTGGGTT) targeting Llph exon 2 were designed using Benchling, and sgRNA synthesized using the Gene Art Precision gRNA Synthesis Kit (Thermo Fisher, A29377) according to kit instructions. In all, 3.43 µg of sgRNA mix was complexed with 40 pmol of Cas9-NLS protein (UC Berkeley Macrolab) for 10 minutes at room temperature. The Cas9-sgRNA RNP along with 0.3 µg of pCE-mp53DD (Addgene #41856, to promote cell survival) were then nucleofected into 4 x 105 hESCs using the P3 Primary Cell 4D-Nucleofector™ X Kit S (Lonza, V4XP-3032) with program CB-150 following manufacturer’s instructions. Cells were then plated in 1 well of a 6-well plate post nucleofection and cultured until confluent to split. Clonal selection was then performed by sparse seeding of 3000 cells in a 10-cm dish, followed by manual picking of clones into individual wells of a 96-well plate.

### Lentivirus production (related to Figure 3)

Replication incompetent lentiviruses were produced in COS1cells by cotransfection of packaging plasmids pMD2.G (Addgene 12259) and psPAX2 (Addgene 12260) with lentiviral vectors: 1) FUW-TetO-Ngn2-T2A-Bsd, where TetO promoter drives expression of full length mouse Neurogenin2 (Ngn2) and Blasticidin S deaminase (Bsd) via cleavage of T2A peptide sequence; 2) FUW-rtTA, to allow for dox-inducible expression of Ngn2-T2A-Bsd construct that is under TetO promoter^107^.

In brief, for each 15-cm plate, 1.05 × 10^7^ of COS1 cells were seeded one day before transfection. 1.9 µg of pMD2.G and 5.63 µg of psPAX2 were mixed with 2.24 µg of each lentiviral vector in 1.5 mL Opti-MEM (Thermo Fisher, 31985070) and 75 µL of 1x polyethylenimine (Sigma-Aldrich, 764965-1G) for 20 minutes at room temperature. Afterwards, plasmid mix was added dropwise onto the plate. Media was replaced the next day with 17 mL of fresh prewarmed DMEM + 10% FBS without Pen/Strep. First collection was done 24 hrs after media replacement, and the virus-containing media was stored in 4 °C. Second collection was done 48 hrs after media replacement, and virus-containing media was combined together. Cell debris was removed by centrifuging at 1,000 ×g for 10 minutes at 4 °C. Lenti X concentrator (Takara, 631232) was then used to concentrate the virus 100x following the manufacturer’s instructions.

### Generation of human induced neurons from H1-hESC with Ngn2 overexpression (related to Figure 3)

Prior to plating cells for induced neurons (iN) differentiation, a 6-well plate was coated with 1:200 Matrigel (Corning, 356234) for at least 2 hours. On day 0, approximately 500k of H1-hESCs are plated per well of 6-well plate in 1mL of mTeSR1 + 2 µM thiazovivin + 2 µg/mL polybrene (Tocris, 7711). 2 µL each of Ngn2-Bsd and rtTA lentivirus were added per well. On day 1, after approximately 18 hours of infection, media was replaced to 2 mL of N3 media (DMEM/F12 1:1 (Gibco, 11320-033) + 1x N2 supplement (Gibco, 17502-048) + 1x non-essential amino-acids (Millipore, TMS-001-C), 5 mg of insulin (dissolved in 10 mM NaOH) (Sigma-Aldrich, I6634-100MG), 0.5x Pen/Strep (Gibco, 15140163)) + 2 µg/mL doxycycline (Dox) (Fisher Scientific, BP26535). On day 2 and 3, media was replaced to 2 mL of N3 media + 2 µg/mL Dox + 2 µg/mL puromycin (Sigma-Aldrich, P8833-25MG). On day 4, media was replaced to 2 mL of N3 media + 2 µg/mL Dox + 2 µg/mL puromycin + 4 µM Cytosine β-D-arabinofuranoside (AraC) (Sigma-Aldrich, C1768).

To prepare iN for imaging, on day 4 glass coverslips were first placed in a 24-well plate, sterilized by washing 3x with 70% ethanol for 5 minutes each, and then UV-irradiated for 30 minutes as the ethanol evaporated. Afterwards, the coverslips were coated with 1:200 matrigel overnight. On day 5 morning, approximately 50k mouse glial cells from newborn wild-type CD1 mice were plated on each coverslip with NBP media (Neurobasal (Gibco, 21103-049), 1x Glutamax (Gibco, 35050061), 1x Gem21 NeuroPlex™ Serum-Free Supplement (Gemini, 400-160), 0.5x Pen/Strep (Gibco, 15140163), 5% iN grade serum - Cytiva HyClone™ (Cytiva, SH30396.03)). On day 5 evening, premature iN cells were detached gently with Accutase, and approximately 150k cells per well were re-plated with 800 µL NBP media + 2 µg/mL doxycycline. On day 7, wash gently with pre-warmed NB zero media (NBP without 5% serum) to remove any dead cells, and replace media with 800 µL NB-2% (NBP with 2% serum instead) + 2 µg/mL Dox + 4 µM AraC. On day 10, partially replace spent media with 300 µL NB-2% + 2 µg/mL Dox + 4 µM AraC. On day 14, stop adding Dox and AraC, and partially replace spent media with 300 µL NB-2%. On day 21, partially replace spent media again with fresh 300 µL NB-2%.

### Human induced neurons morphology analysis (related to Figures 3C and 3D)

Glia co-cultured LLPH^+/+^ and LLPH^Nterm/Nterm^ hiNs were grown until DIV 30 as mentioned above. Cells on the glass coverslips were washed carefully with cold PBS (Fisher Scientific, BP2944100) once and then fixed in 4% paraformaldehyde (Fisher Scientific, 43368-9M), 4% sucrose (Sigma-Aldrich, 8510-500GM), in PBS for 20 minutes at 4 °C. Afterwards, the coverslips were washed three times with room temperature PBS and blocked in 2.5% goat serum (MP Biochemical, 092939249), 2.5% BSA (Sigma-Aldrich, A7906-100G), and 0.2% Triton X-100 (Sigma-Aldrich, X100-500ML) in PBS for 1 hr at room temperature. Primary antibody MAP2 (chicken, 1:1000, Encor, CPCA-MAP2) was added in the same blocking buffer and left to incubate overnight at 4 °C. Coverslips were then washed in PBS three times and incubated with fluorescence-labeled secondary antibodies (goat anti chicken Alexa 647, 1:1500, Invitrogen, A-21449) in PBS for 1 hour at room temperature. Coverslips were again washed in PBS three times and then incubated with (4’,6-diamidino-2-phenylindole) (DAPI; Thermo Fisher, 62248) diluted 1:10,000 in PBS. Afterwards, slides were mounted on microscope slides with Fluoromount-G (SouthernBiotech, 0100-01). Images were taken by Zeiss LSM 780 and analyzed by SNT plugin in Fiji^108^.

### Ribosome profiling on human induced neurons (related to Figures 3E-G)

LLPH^+/+^ and LLPH^Nterm/Nterm^ hiNs were grown until DIV 14 as mentioned above, with modifications as stated below. Glia was not added on day 5 and premature iN cells were not detached and instead left growing on matrigel coated 6-well plate for the full 14 days. On day 5, media was fully replaced to NB zero + 2 µg/mL Dox + 4 µM AraC. On day 10, 50% of spent media was replaced with NB zero + 2 µg/mL Dox + 4 µM AraC. hiNs were harvested using Accutase in the presence of 100 µg/mL CHX and lysed with lysis buffer B as mentioned above (Sucrose Gradient Fractionation for mouse E12.5 tissues). Lysis was performed and the cytoplasmic fraction was enriched by serial centrifugation as described previously (RAPIDASH for E14 mouse embryonic stem cells).

Ribosome profiling libraries were prepared by generally following the published protocol of McGlincy and Ingolia^109^ with modifications below. To isolate RNA input samples for RNA-seq, 10 µL of cytoplasmic lysate is transferred into 500 µL of cold TRIzol, mixed by pipetting, and stored at −80 °C for subsequent RNA extraction using NEBNext® Ultra II Directional RNA Library Prep Kit for Illumina (NEB, E7760L). The remaining lysate (∼375 µg total RNA) was treated with 0.33 µg RNase A (Thermo Scientific, AM2271) and 200 U RNase T1 (Thermo Scientific, EN0541) for 30 min on rotator at room temperature. Digestion was stopped by adding 10 µL SUPERaseIn and transferring the samples on ice. Meanwhile, to isolate intact 80S ribosomes and ribosome footprints (RFPs), 10-50% linear sucrose gradients were made in 14×89 mm open-top polyallomer centrifuge tubes (Beckman Coulter, 331372) using a gradient maker (Biocomp, 108). Before layering the lysates onto the gradient, 200 μL sucrose gradient solution was removed from each centrifuge tube. 200 μL cytoplasmic lysate containing 20 μg RNA was used for each RFP sample. The tubes were centrifuged using a SW 41 Ti swinging bucket rotor (Beckman Coulter, 331336) at 40,000 rpm for 2.5 hours at 4 °C.

Fractions were collected for 30 seconds each using the Density Gradient Fraction System with a flow rate of 1.5 mL/minute (Brandel, BR-188l). 80S were found in fraction 7 for hiN D14 LLPH^+/+^ rep 2 and LLPH^Nterm/Nterm^ rep 1, and fraction 6 and 7 for the other samples. A 3x volume of TRIzol LS (Thermo Fisher, 10296028) was added to these fractions. RFP samples were then extracted from TRIzol using the Direct-Zol Microprep Kit (Zymo, R2060) according to the manufacturer protocol. Subsequently, these samples were treated with TurboDNAse (Thermo Fisher, AM2238) at 37 °C for 30 minutes according to the manufacturer protocol and cleaned up with Zymo RNA Clean & Concentrator Kit (Zymo, R1013) with some modifications to enrich for small RNAs (>17 nt, <200 nt). Adjusted binding buffer was prepared by mixing equal volumes of 100% ethanol (Gold Shield, 412804) and RNA binding buffer. Samples were first adjusted to 100 µL with nuclease free water (Thermo Fisher, 10977023). Then, 200 µL of adjusted binding buffer was added and mixed, followed by adding 450 µL of 100% ethanol and mixed. Samples were then processed with spin columns as per manufacturer protocol and eluted twice with 6 µL of nuclease free water each.

RFPs were denatured at 80°C for 90 s in denaturing sample loading buffer (10 mM EDTA and bromophenol blue (Fisher Scientific, MBX14107), in formamide (Promega, H5052)), then incubated on ice for 5 min before running on a 15% Tris-borate-EDTA-urea (TBE-urea) polyacrylamide gel. Fragments were size-selected using NI-800 and NI-801^109^ as 26–34 nt markers. Gel slices were freeze-thawed for at least 30 min at −80°C and 3 minutes at room temperature. They were then crushed by forcing the gel through a hole in 0.5 mL tube pierced by 20 gauge needle (Santa Cruz Biotechnology, sc-359535) by centrifuging at 15,000 ×g for 2 minutes. RNA were then extracted from the minced gel at room temperature overnight in 400 µL RNA extraction buffer^109^, then re-extracted with an additional 200 µL RNA extraction buffer for additional 3 hours, removing the minced gel by using 0.22 µm cellulose acetate centrifuge tube filters (Corning, 8160) and spinning at max speed for 2 minutes at room temperature. The combined 600 µL extraction was precipitated with 2 µL GlycoBlue (Thermo Fisher, AM9516) and 750 µL 100% isopropanol (Sigma-Aldrich, PX1835-6) overnight at −80°C. Precipitated RFPs were pelleted at 21,000 ×g for 30 min at 4°C, washed with ice-cold 80% ethanol in water, air dried at room temperature for 10 min, then dissolved in 4 µL of 10 mM Tris pH 8, dephosphorylated, and ligated to barcoded linkers as per the published protocol. RNA linkers were pre-adenylated beforehand using Mth RNA ligase (NEB, E2610S). Unreacted linker was deadenylated and digested as per the published protocol. The barcoded RFPS were then purified using Zymo Oligo Clean & Concentrator column (Zymo, D4060) according to the manufacturer’s protocol, and subsequently reverse-transcribed as per the published protocol. Template RNA was degraded by alkaline hydrolysis, and cDNA was purified using Zymo Oligo Clean & Concentrator column, denatured in sample loading buffer, and size selected on a 10% TBE-urea gel, as marked by NI-800 and NI-801 that had been processed in parallel with the samples. Gel slices were processed as described previously, and cDNA was extracted at room temperature overnight in 400 µL DNA extraction buffer^109^, then re-extracted with an additional 200 µL DNA extraction buffer for 3 hrs. The cDNA was pelleted, washed, and dried as described above, then resuspended in 15 µL 10 mM Tris pH 8. cDNA was circularized by adding 2 µL 10X CircLigase I buffer, 1 µL of 1 mM ATP, 1 µL of 50 mM MnCl2, and 1 µL of CircLigase I (Lucigen CL4111K) to the 15 µL of cDNA and incubating at 60°C for 12 hr, then 80°C for 10 min. Using 1 µL of circularized DNA template per 50 µL reaction, library construction to add indexing primers was performed as per the published protocol using a different reverse primer for each replicate. PCR products were purified using a Zymo DNA Clean & Concentrator column (Zymo, D4003). Size selection was performed on a 8% TBE-urea gel, with the lower bound marked by NI-803^109^ that had undergone library construction in parallel with the samples, and the upper bound at 170 nt as marked by O’Range 20 bp DNA ladder (Thermo, SM1323). Gel slices were extracted as described above. DNA was precipitated as described above, except using 1.25 µL of 20 µg/mL glycogen (Themo Fisher, 10814-010) instead of GlycoBlue. The pellet was resuspended in 15 µL of 10 mM Tris pH 8. Library quality was analyzed on an Agilent 2100 Bioanalyzer (High-Sensitivity DNA) at the Stanford Protein and Nucleic Acid Facility. Library concentration was measured using Qubit dsDNA high sensitivity kit (Thermo Fisher, Q33231), and the RFP samples were then pooled in equal amounts.

RNA samples were mixed together with RFP samples in 4:6 ratio. Libraries were sequenced by Novogene (Sacramento, CA) on a full lane Illumina NovaSeq X Plus with paired-end 150 bp reads.

### Ribosome profiling analysis (related to Figures 3E-G)

Due to the short insert length, only analysis of Read 1 was necessary. cutadapt version 2.4^110^ was used to trim 3′ adapter sequences from Read 1 with parameters “-j 0 -u 3 -a AGATCGGAAGAGCACAGTCTGAACTCCAGTCAC --discard-untrimmed -m 15”. In-line barcodes were demultiplexed using fastx_barcode_splitter.pl (http://hannonlab.cshl.edu/fastx_toolkit/) with parameters “--eol”. Unique molecular identifiers and in-line barcodes were extracted using umi_tools version 1.0.1^111^ with parameters “extract -- extract-method=string --bc-pattern=NNNNNCCCCC -–3prime”. Reads were filtered by quality using fastq_quality_filter (http://hannonlab.cshl.edu/fastx_toolkit/) with parameters “-Q33 -q 20 -p 70 -z”. To remove reads originating from rRNA, transfer RNA (tRNA), and small nuclear RNA (snRNA), reads aligning to these sequences using bowtie2 version 2.3.4.3^112^ with parameters “-L 18” were discarded. PCR duplicates were then removed using UMI-tools. Ribosome A site positions were determined by offsetting the distance of the 5’ end of each RFP read to canonical start sites in each length group and adding 4 nucleotides. Reads aligning to the CDS were used for RFP libraries, and reads aligning to the entire transcript were used for RNA-Seq libraries. Reads mapping to mitochondrial DNA genes were excluded from further analysis. Transcripts with counts per million (CPM) >2 were retained for downstream analysis RFP and RNA-seq libraries were normalized separately by the trimmed mean of M-values method in edgeR^113^. Differential RFP and RNA-seq abundance and enrichment were analyzed using voom^114^ and limma^115^. Multiple testing correction was performed using the Benjamini-Hochberg method.

### RAPIDASH on macrophages (related to Figures 5A and 5B)

RAPIDASH isolation of ribosome complexes from macrophages was performed as described above with the following exceptions. For each sample, ∼100 million macrophages were used as starting material. For CHX treatment of macrophages, 100 µg/mL CHX was added to the BMDM culture media for seven minutes before cells were washed twice in ice cold PBS containing 100 µg/mL CHX and harvested with a cell scraper.

### Proteomic analysis of ribosomal complexes in macrophages (related to Figures 5A and 5B)

#### In solution digestion

Aliquots of macrophage ribosomal samples were digested with trypsin for proteomic analysis. Samples (3 µg protein for label free experiments, 30 µg protein for Tandem Mass Tags (TMT) experiments) were prepared at a concentration of 1 mg/ml in a solution containing 8M Guanidinium Hydrochloride (GndHCl) and 50 mM ammonium bicarbonate and frozen till processing. After thawing, samples were added 10 mM tris(2-carboxyethyl)phosphine and incubated for 1h at 56°C. This was followed by addition of 20 mM iodoacetamide and a 30-minute incubation at room temperature in the dark. The samples were then diluted with 20 mM ammonium bicarbonate to reach a concentration of 1M GndHCl. For digestion, the samples were then added 3% (W/W) Lysyl Endopeptidase®, Mass Spectrometry Grade (Lys-C) (FUJIFILM Wako Chemicals U.S.A. Corporation) and incubated at 37°C on a shaker overnight. After that, samples were added to 3% (W/W) trypsin MS grade (Pierce-Thermo Fisher) and digested for 6 additional hours. After this, samples were acidified with formic acid to a final concentration of 5% formic, and the digests were then desalted using either C18 ZipTip (Millipore) or 100 μL OMIX C18 pipette tips (Agilent) following the manufacturer’s protocol. Eluates were dry-evaporated in preparation for direct label free MS analysis or for labeling with tandem mass tag (TMT) reagents for quantitative comparison.

#### TMT labeling

Dried samples were labeled according to TMTProTM-18 label plex kit instructions (Thermo Fisher Scientific), with some modifications. Briefly, samples were resuspended in 16 μL 0.1M triethylammonium bicarbonate pH 8.0. TMT reagents were dissolved in acetonitrile at 25 µg/μL, and 5 μL of these stocks added to the samples. After incubation for 1 h at room temperature samples were quenched with 1 µl 5% hydroxylamine, then all 15 samples combined and partially evaporated in speedvac until volume was around 5 µl. 100 μL 1% formic acid was added and then peptides desalted using a C18 SepPak. The SepPak eluate was dried in preparation for fractionation by high pH reverse phase chromatography.

#### High pH reverse phase chromatography

SepPak cleaned,TMT labeled samples were resuspended in 240 µl 20mM ammonium formate pH 10.4 for fractionation of the peptide mixture by high pH RP chromatography using a Phenomenex Gemini 5u C18 110A 150 x 4.60 mm column, operating at a flow rate of 0.550 mL/min. Buffer A consisted of 20 mM ammonium formate (pH 10), and buffer B consisted of 20 mM ammonium formate in 90 acetonitrile (pH 10). Gradient details were as follows: 1 to 30 B in 49 min, 30 B to 70 B in 4 min, 70 B down to 1 B in 4 min. Peptide-containing fractions were collected, evaporated, and resuspended in 0.1 formic acid for mass spectrometry analysis.

#### Mass spectrometry analysis

Peptide digests resuspended in 0.1% formic acid were injected (approximately 2 µg) onto a 2 µm 75 µm x 50 cm PepMap RSLC C18 EasySpray column (Thermo Scientific). For peptide elution, 3-hour water/acetonitrile gradients (2–25% in 0.1% formic acid) were used, at a flow rate of 200 nl/min. Analysis of the label free samples was performed in an Orbitrap Lumos Fusion (Thermo Scientific) in positive ion mode. MS spectra were acquired between 375 and 1500 m/z with a resolution of 120000. For each MS spectrum, multiply charged ions over the selected threshold (2E4) were selected for MSMS in cycles of 3 seconds with an isolation window of 1.6 m/z. Precursor ions were fragmented by HCD using a relative collision energy of 30. MSMS spectra were acquired in centroid mode with resolution 30000 from m/z=110. A dynamic exclusion window was applied which prevented the same m/z from being selected for 30s after its acquisition.

For analysis of the TMT experiments, aliquots of 10 non-consecutive chromatographic fractions were analyzed in an Orbitrap Exploris 480 (Thermo Scientific) in positive ion mode. MS spectra were acquired between 375 and 1500 m/z with a resolution of 120000. For each MS spectrum, multiply charged ions over the selected threshold (2E4) were selected for MS/MS in cycles of 3 seconds with an isolation window of 0.7 m/z. Precursor ions were fragmented by HCD using stepped relative collision energies of 30, 35 and 40 to ensure efficient generation of sequence ions as well as TMT reporter ions. MS/MS spectra were acquired in centroid mode with resolution 60000 from m/z=120. A dynamic exclusion window was applied which prevented the same m/z from being selected for 30s after its acquisition.

#### Peptide and protein identification and quantitation

Peak lists were generated using PAVA in-house software^116^. All generated peak lists were searched against the mouse subset of the SwissProt database (SwissProt.2019.07.31, 17026 entries searched), using Protein Prospector^117^ with the following parameters: Enzyme specificity was set as Trypsin, and up to 2 missed cleavages per peptide were allowed. Carbamidomethylation of cysteine residues, and, in the case of TMT labelled samples, TMT16plex labeling of lysine residues and N-terminus of the protein, were allowed as fixed modifications. N-acetylation of the N-terminus of the protein, loss of protein N-terminal methionine, pyroglutamate formation from peptide N-terminal glutamines, oxidation of methionine were allowed as variable modifications. Mass tolerance was 10 ppm in MS and 30 ppm in MS/MS. The false positive rate was estimated by searching the data using a concatenated database which contains the original SwissProt database, as well as a version of each original entry where the sequence has been randomized. A 1% FDR was permitted at the protein and peptide level. For quantitation only unique peptides were considered; peptides common to several proteins were not used for quantitative analysis. For TMT based quantitation, relative quantization of peptide abundance was performed via calculation of the intensity of reporter ions corresponding to the different TMT labels, present in MS/MS spectra. Intensities were determined by Protein Prospector. Median intensities of the reporter ions (each TMT channel) for all peptide spectral matches (PSMs) were used to normalize individual (sample specific) intensity values. For each PSM, relative abundances were calculated as ratios vs the average intensity levels in the 3 channels corresponding to control (non-stimulated) samples. For total protein relative levels, peptide ratios were aggregated to the protein levels using median values of the log2 ratios. Statistical significance was calculated with a 2-tailed t-test.

### Sucrose gradient fractionation experiments for macrophages (related to Figures 5C-5F)

For Puromycin treatment of BMDMs, Puromycin (2 mM, Sigma) was added to the BMDM culture media for 10 minutes before cells were washed twice in ice cold PBS and harvested with a cell scraper. Cell pellets were lysed immediately or frozen and stored at −80°C.

For polysome profiling experiments, 30-40 million macrophages were lysed in one of two polysome buffers: buffer A (20uM HEPES-KOH pH 7.6, 15mM magnesium acetate, 60mM ammonium chloride, 1mM DTT, 1% Triton-X, 0.5% sodium deoxycholate, 8% glycerol, 0.02 U/uL TURBO DNAse, 1x Halt Protease Inhibitor Cocktail) or buffer B (20mM Tris pH 7.4, 150mM NaCl, 15mM MgCl_2_, 1mM DTT, 0.5% Triton-X, 8% glycerol, 0.02 U/uL TURBO DNase, 1x Halt Protease Inhibitor Cocktail). Lysates were vortexed three times for 30 seconds, then incubated on ice for 30 minutes. Macrophages lysed in buffer A were centrifuged at 800 ×g for 5 minutes, then 8000 ×g for 5 minutes, then at max speed for 5 minutes in a tabletop microcentrifuge at 4°C. Macrophages lysed in buffer B were centrifuged at 1,800 ×g for 5 minutes, then at 10,000 ×g for ten minutes in a tabletop microcentrifuge at 4°C.

Puromycin-treated macrophages were lysed in buffer A supplemented with 2 mM Puromycin and 0.2 U/µL SUPERase-IN RNase Inhibitor. Non-Puromycin-treated (control) macrophages were lysed in buffer A supplemented with 100ug/mL CHX and 0.2 U/µL SUPERase-IN RNase Inhibitor.

Puromycin and non-Puromycin-treated macrophage lysates (300 µL) were loaded onto a 10-50% sucrose gradient (20 mM Tris pH 7.4, 100 mM KCl, 4 mM MgCl_2_) generated using a Gradient Master 108 (Biocomp).

For RNase treatment of macrophage samples, macrophages were treated *in vitro* with CHX and lysed in polysome buffer B supplemented with 100 µg/mL CHX. Control samples were lysed in buffer B supplemented with 100 µg/mL CHX and 0.2 U/µL SUPERase-IN RNase Inhibitor. Following centrifugation, the RNA concentration in each sample was measured using a Nanodrop 2000c Spectrophotometer (Thermo Scientific). For every 75 µg of RNA, 0.5 µL RNase A (AM2271, Ambion) and 0.3 µL RNase T1 (EN0541, Thermo Scientific) were added, and samples were incubated with rotation at room temperature for 30 minutes (control samples were meanwhile kept on ice.) After 30 minutes, SUPERase-In RNase Inhibitor was added to the RNase-treated samples, such that the volume of SUPERase-IN added was equal to twice the volume of the combined RNases that were added. RNase-treated and control macrophage lysates were loaded onto a 10-50% sucrose gradient (20 mM Tris pH 7.4, 150 mM NaCl, 15 mM MgCl_2_).

All samples were centrifuged at 38,000 rpm for 2.5 hours at 4°C in a Beckman L8-70M ultracentrifuge. Samples were separated into 14 fractions on a Piston Gradient Fractionator (Biocomp).

### Western blotting for macrophage gradient fractions (related to Figures 5C-F)

Protein was precipitated from polysome fractions using the ProteoExtract Protein Precipitation kit (Sigma-Aldrich) according to the manufacturer’s instructions with the following modifications: 700 µL of Precipitation Agent was added to 700 µL of each polysome fraction, and samples were incubated for at least 24 hours at −20°C. All centrifugation steps were performed at max speed at room temperature in a tabletop microcentrifuge (Eppendorf).

Precipitated protein was resuspended in 35 µL of 1x Laemmli Sample Buffer supplemented with 10% 2-mercaptoethanol. 10uL of each sample was loaded into 4-15% Mini-PROTEAN TGX Precast protein gels and transferred onto nitrocellulose membranes. Membranes were blocked in TBS-T with 5% milk and incubated overnight in TBS-T with 5% milk at 4°C with the following primary antibodies: anti-RPL23 (A305-008A, Bethyl), anti-RPS6 (2317S, Cell Signaling), anti-RPS12 (16490-1-AP, Proteintech), anti-Viperin (MaP.VIP, Abcam), anti-CMPK2 (H00129607-A01, Abnova), anti-HO-1 (ADI-SPA-895-F, Enzo), anti-RACK1 (4716S, Cell Signaling), and anti-OAS3 (21915-1-AP, Proteintech). Membranes were washed and incubated for one hour at room temperature with either Promega Anti-Mouse (W402B) or Anti-Rabbit (W401B) IgG HRP Conjugates. Membranes were incubated for two minutes with SuperSignal West Dura Extended Duration Substrate (Protein Biology) and visualized using the BioRad ChemiDoc Touch Imaging System.

### LLPH-Flag immunoprecipitation (related to Supplementary Figure 2A)

A549 cells were transduced with lentiviral particles expressing LLPH-Flag or GFP-Flag under the control of inducible TRE3G promoter (TRE3G-LLPH-Flag or TRE3G-GFP-Flag) and Tet3G under constitutively expressed PGK promoter (PGK-Tet3G-IRES-mCherry). Cells stably expressing mCherry were screened to generate A549 cell lines with inducible LLPH-Flag or GFP-Flag. Cells were treated with 1 µg/mL Dox for 48 hours to induce LLPH-Flag or GFP-Flag expression. Cells were then harvested with CHX treatment as described previously (RAPIDASH for E14 mouse embryonic stem cells - Harvest and cytoplasmic lysis). Mock pull downs prepared in lysates expressing GFP-flag were performed in parallel as controls for non-specific binding. To prepare cytoplasmic lysate, cells were lysed with cold <u>lysis buffer C</u> (20 mM HEPES-KOH pH 7.5, 150 mM potassium acetate (KOAc), 5 mM MgCl_2_, 2 mM DTT, 8% glycerol, 1% Triton X-100, 0.5% sodium deoxycholate, 100 μg/ml CHX, 100 U/ml SUPERase In, 25 U/ml TurboDNase, and 1× Halt™ Protease and Phosphatase Inhibitor) by vortexing at high speed for 30 seconds and putting them back on ice for 30 seconds. This was repeated another two times for a total of 3 minutes. Afterwards, samples were incubated for 30 minutes on ice with vortexing every 10 minutes. Lysates were cleared by sequential centrifugation at 800 ×g for 5 minutes, 800 ×g for 5 minutes, 8000 ×g for 5 minutes and 21,300 ×g for 10 minutes at 4 °C. Cytoplasmic lysates were then treated with 1 µL RNase T1 and 1 µL MNase (Thermo Fisher, EN0181) for 30 minutes at room temperature. The reaction was quenched by addition of 5 µL of SUPERaseIn and leaving the samples on ice. To perform the immunoprecipitation, 300 µL of cytoplasmic lysate was incubated with 50 µL of ANTI-FLAG® M2 Affinity Gel (Sigma-Aldrich, A2220) for 2 hours on a turning wheel at 4 °C. Beads were then transferred into 15 mL falcon tubes and batch washed two times with 10 mL wash buffer A (20 mM HEPES-KOH pH 7.5, 150 mM KOAc, 5 mM MgCl_2_, 1 mM DTT, 2% glycerol, 0.01% NP40 (Thermo Fisher, 85124)) each time and then batch washed once with 10 mL of buffer B (20 mM HEPES-KOH pH 7.5, 150 mM KOAc, 5 mM MgCl2, 1 mM DTT, 2% glycerol, 0.05% octaethylene glycol monododecyl ether (Nikkol; Sigma-Aldrich, P8925)). Afterwards, 5 mL buffer B was added to each sample, and the slurry was transferred to a Mobicol column (Boca Scientific) to perform the last column wash. Proteins were then eluted in 100 µL of 250 µg/mL Flag peptide (Sigma-Aldrich, F3290) in buffer B by incubating the column on a turning wheel at 4 °C for 45 minutes. Proteins were partially separated on SDS-PAGE gels, allowing the bromophenol blue marker to reach 1 cm inside the gel. Gel was stained using ProtoBlue Safe Colloidal Coomassie Blue G-250 stain. The upper portion of the lanes containing the proteins was excised and digested in-gel with trypsin as described previously^118^. The extracted digests were vacuum-evaporated and dried samples were resuspended in 5 ul 0.1% formic acid and subjected to mass spectrometry analysis, as described above (Proteomic analysis of ribosomal complexes in macrophages - Mass spectrometry analysis, peptide and protein identification and quantitation)

### Cell viability assay (related to Supplementary Figure 2D)

hESCs were passaged as described above (H1-hESC cell culture and CRISPR), and cells were counted to ensure even plating across genotypes. Cells were plated in Geltrex-coated black-sided clear-bottomed 96-well tissue culture dishes (Sigma-Aldrich, CLS3603). Wells on the edge of the dish were excluded. Cell viability was measured using the CellTiter-Glo Luminescent Cell Viability Assay (Promega, G9242) following kit instructions. Significance was measured using Student’s t tests.

---

**Figure S1:**
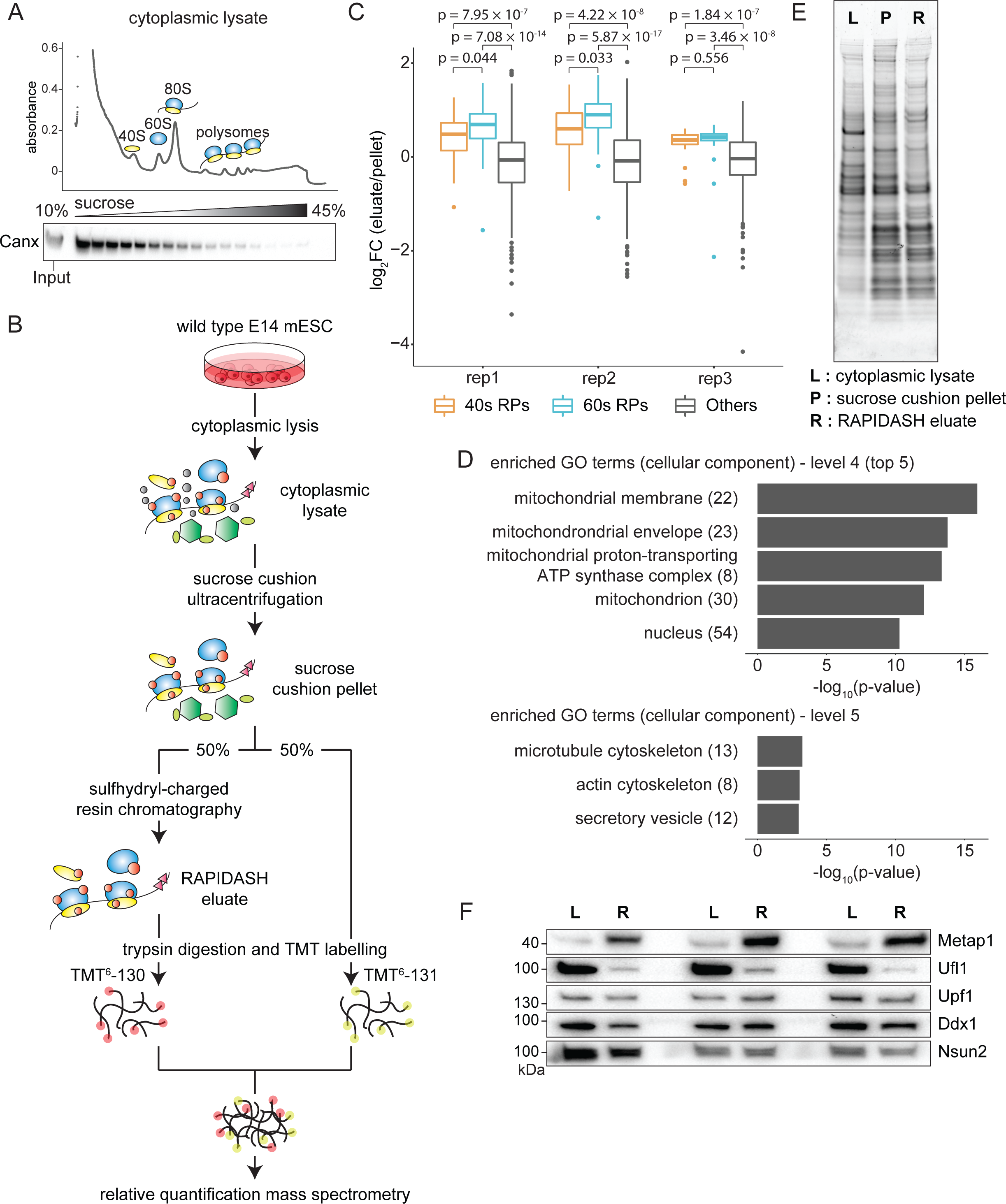
Additional information on RAPIDASH characterization in E14 mouse embryonic stem cells (mESCs). (A) Top: Sucrose gradient fractionation of E14 mESC cytoplasmic lysate as a control for RAPIDASH characterization in Figure 1C. Bottom: Western blot of the fractions probed for Canx, a marker for endoplasmic reticulum (ER) microsomes, shows that ER microsomes have heterogeneous densities and are present throughout most fractions. (B) Schematic of the strategy to compare relative ribosome enrichment between sucrose cushion centrifugation alone and the complete RAPIDASH workflow using TMT mass spectrometry. Wild-type E14 mESCs were subjected to cytoplasmic lysis, with 50% of the material being processed by sucrose cushion ultracentrifugation alone, and another 50% being subjected to an additional sulfhydryl-charged resin chromatography step. The enriched proteins were digested to peptides, which were labeled with TMT reagents to allow for relative quantification by LC-MS/MS. The peptides from each sample were combined and analyzed by mass spectrometry. Three biological replicates were performed. (C) Refer to Figure 1D: more detailed breakdown of the boxplot of normalized log_2_ FC of RAPIDASH eluate over sucrose cushion ultracentrifugation pellet TMT mass spectrometry ratios, with ribosomal proteins (RPs) being further divided into 40S (small subunit) RPs (orange), and 60S (large subunit) RPs (cyan). There is a slight enrichment of 60S RPs over 40S RPs that is significant (p-values < 0.05) in 2 out of 3 biological replicates based on Welch’s t-test. (D) Analysis of GO terms of proteins depleted in RAPIDASH eluate over sucrose cushion pellet TMT mass spectrometry data. Proteins were analyzed by Manteia^30^ for Gene Ontology Cellular Component (GOCC). The top terms in level 4 and level 5 are shown. (E) Refer to Figure 1E: SYPRO Ruby blot stain of E14 mESCs sucrose cushion pellet and RAPIDASH eluate samples. Protein bands at lower molecular weight indicate enrichment of ribosomal proteins in sucrose cushion pellet and RAPIDASH eluate samples relative to cytoplasmic lysate. (F) Refer to Figure 1F: all three biological replicates for western blotting analysis of known RAPs enriched by RAPIDASH (R) compared to lysate.

**Figure S2:**
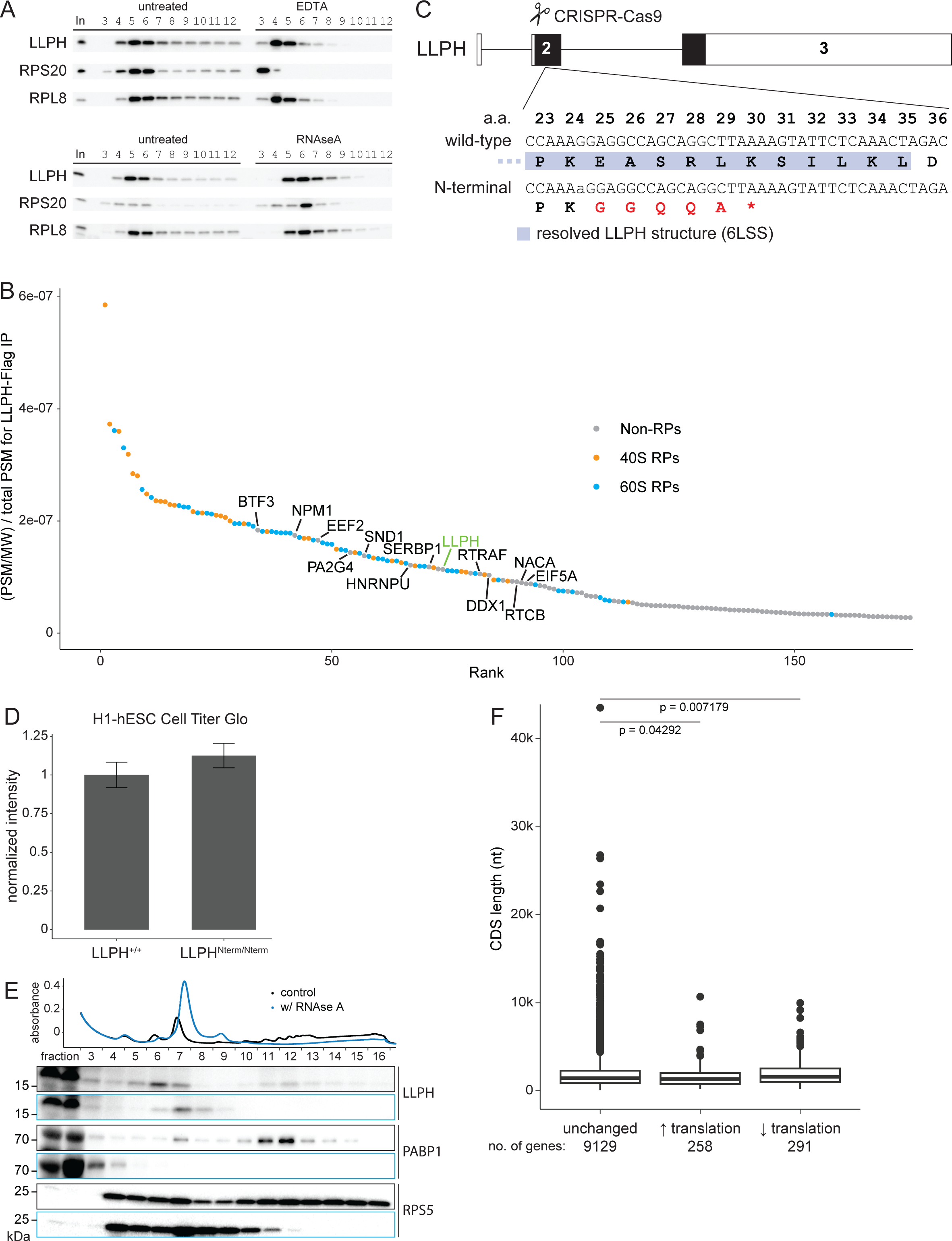
Additional characterization of the LLPH interactome, LLPH CRISPR design, and the hESC and hiN LLPH^Nterm/Nterm^ mutants. (A) Refer to Figure 3B: additional western blot data of sucrose gradient fractionation in P493-6 cells upon treatment with EDTA or RNAseA. Rpl8 and Rps20 were shown as controls for large and small subunits respectively. (B) Rank order plot of normalized peptide spectrum match (PSM) of proteins identified in LLPH-Flag immunoprecipitation (IP) from cytoplasmic lysate. PSM was normalized based on molecular weight (MW) and total PSM detected for each sample. 40S RPs are shown as orange, and 60S RPs are shown as cyan. Given overall high ranks of RPs, LLPH in cytoplasm largely binds with ribosomes. Top non-RPs in LLPH interactome hint at possible role in translational control. (C) Schematic for CRISPR editing of the LLPH gene. CRISPR editing of the endogenous LLPH gene in H1-hESC leads to the introduction of early stop codon, resulting in expression of the N-terminal 24 amino acids plus five extra amino acids. The expressed N-terminal portion of LLPH that is highlighted in blue was resolved in the cryo-EM structure of the human pre-60S particle (PDB ID: 6LSS). (D) Cell viability of LLPH^+/+^ and LLPH^Nterm/Nterm^ H1-hESCs. LLPH^+/+^ and LLPH^Nterm/Nterm^ H1-hESC viability was assessed by performing a CellTiter-Glo assay. (E) Confirmation of LLPH as an RNA-independent RAP in hiNs DIV 5. Sucrose gradient fractionation was performed on hiNs DIV 5 lysate that was treated with (blue) or without (black) RNase A. Proteins in each fraction were precipitated and analyzed by western blotting for the presence of LLPH. Rps5 is shown as a control for an RP. Pabp1 is shown as a control for an RNA-binding protein that cofractionates with the ribosome. (F) The complete data used in the analysis for Figure 3G. Genes downregulated for translation tend to have longer coding sequences (CDSs) than those that are translationally unchanged (Mann-Whitney U test, p = 0.0072).

**Figure S3:**
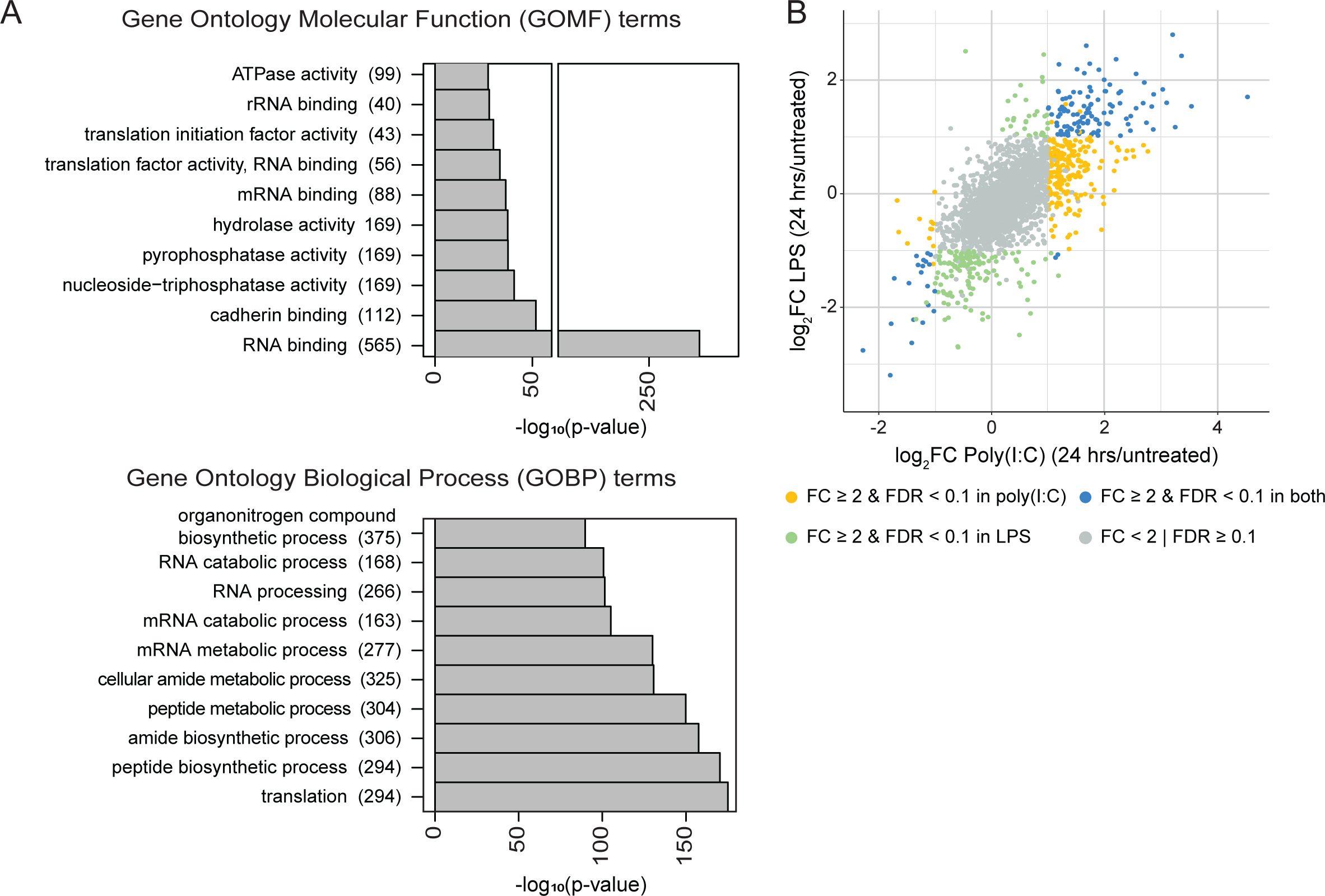
Additional information on novel RAPs in macrophages following TLR activation. (A) Analysis of GO terms in unstimulated BMDMs MS data with three biological replicates. Proteins detected in all three replicates were analyzed by Manteia^30^ for GOMF and GOBP terms level 4 or higher. Top 10 GO terms are shown. (B) Comparison of proteins enriched by RAPIDASH 24 hours after LPS vs. poly(I:C) stimulation of BMDMs. Proteins that pass the cutoffs (FC ≥ 2 & FDR < 0.1) only in LPS-stimulated BMDMs are green, those that pass the cutoffs only in poly(I:C)-stimulated BMDMs are yellow, and those that pass the cutoffs in both LPS and poly(I:C)-stimulated BMDMs are blue. Proteins that do not pass any cutoff are gray.

